# Assessment of differentially methylated loci in individuals with end-stage kidney disease attributed to diabetic kidney disease

**DOI:** 10.1101/2020.07.30.228734

**Authors:** LJ Smyth, J Kilner, V Nair, H Liu, E Brennan, K Kerr, N Sandholm, J Cole, E Dahlström, A Syreeni, RM Salem, RG Nelson, HC Looker, C Wooster, K Anderson, GJ McKay, F Kee, I Young, NICOLA Collaborative Team, Warren 3 and Genetics of Kidneys in Diabetes (GoKinD) Study Group, D Andrews, C Forsblom, JN Hirschhorn, C Godson, PH Groop, AP Maxwell, K Susztak, M Kretzler, JC Florez, AJ McKnight, on behalf of the GENIE consortium

## Abstract

A subset of individuals with type 1 diabetes mellitus (T1DM) are predisposed to developing diabetic kidney disease (DKD), which is the most common cause globally of end-stage kidney disease (ESKD). Emerging evidence suggests epigenetic changes in DNA methylation may have a causal role in both T1DM and DKD. The aim of this investigation was to assess differences in blood-derived DNA methylation patterns between individuals with T1DM-ESKD and individuals with long-duration T1DM but no evidence of kidney disease upon repeated testing. Blood-derived DNA from individuals (107 cases, 253 controls and 14 experimental controls) were bisulphite treated before DNA methylation patterns from both groups were generated and analysed using Illumina’s Infinium MethylationEPIC BeadChip arrays (n=862,927 sites). Differentially methylated CpG sites (dmCpGs) were identified (false discovery rate adjusted p≤×10^−8^ and fold change ±2) by comparing methylation levels between ESKD cases and T1DM controls at single site resolution. Gene annotation and functionality was investigated to enrich and rank methylated regions associated with ESKD in T1DM.

Top-ranked genes within which several dmCpGs were located and supported by *in silico* functional data, and replication where possible, include; *AFF3, ARID5B, CUX1, ELMO1*, *FKBP5*, *HDAC4, ITGAL, LY9*, *PIM1, RUNX3, SEPTIN9*, and *UPF3A*. Top-ranked enrichment pathways included pathways in cancer, TGF-β signalling and Th17 cell differentiation.

Epigenetic alterations provide a dynamic link between an individual’s genetic background and their environmental exposures. This robust evaluation of DNA methylation in carefully phenotyped individuals, has identified biomarkers associated with ESKD, revealing several genes and implicated key pathways associated with ESKD in individuals with T1DM.

## Main Text Introduction

Type 1 diabetes mellitus (T1DM) is a polygenic disease characterised by autoimmune destruction of the insulin producing beta cells in the pancreas which subsequently leads to hyperglycaemia (1). Up to 40% of individuals with T1DM are predisposed to diabetic kidney disease (DKD), a microvascular diabetic complication (2,3). Diabetes remains the primary disease causing end-stage kidney disease (ESKD) (4,5). Individuals with DKD are also at increased risk of developing cardiovascular disease and premature mortality (6).

Emerging evidence indicates that autoimmune diseases such as T1DM are influenced by the interaction of genetic and epigenetic factors (7). Genome-wide association studies (GWAS) have identified >50 loci associated with T1DM risk (8,9), including rs3802604 in *GATA3*, rs1770 in *MHC* (10), rs2292239 in *ERBB3* (11) and rs2476601 in *PTPN22* (12).

GWAS have been completed in populations with T1DM assessing the genetic predisposition to DKD and ESKD. One recent GWAS encompassing almost 20,000 individuals with T1DM (13) identified 16 risk loci including the missense mutation rs55703767 in *COL4A3*. Additional GWAS have identified nucleotide variants in rs7583877 in *AFF3* (14), rs1888747 and rs10868025 in *FRMD3*, (15) and rs4972593 in *CDCA7-SP3* (16).

Epigenetic changes in DNA methylation have been associated with T1DM (17) and DKD (18) using peripheral whole blood samples. Bell et al. (19) used Illumina’s Infinium 27K array in 2010 to assess epigenetic profiles associated with DKD in T1DM within 192 individuals. Nineteen differently methylated CpG sites (dmCpGs) were reported and correlated with time to the development of DKD. Six genes of interest with at least two dmCpGs including *CUX1*, *ELMO1*, *FKBP5*, *PRKAG2* and *PTPRN2* were identified from an epigenome-wide association study (EWAS) published in 2014 (20). This investigation included individuals with DKD, caused by either T1DM or type 2 diabetes mellitus (T2DM).

In 2014, Gu et al. (21) employed bisulphite pyrosequencing of 778 individuals with T1DM, with and without DKD, and reported a decrease in the DNA methylation levels within *IGFBP1.* In 2015, Swan et al. (22) assessed DNA methylation variation in genes which encode mitochondrial proteins using Illumina’s 450K and 27K methylation arrays in 442 individuals with T1DM and DKD. In all, 46 dmCpGs were identified in both individuals with DKD and ESKD. The largest change in methylation was evident for cg03169527 within *TAMM41*. Previous EWAS have identified differential methylation features with associations to chronic kidney disease (CKD) (23–26). Improvements in sequencing and profiling technologies have prompted a rise in the number of studies assessing diseases using these techniques for biomarker discovery (23,27).

The aim of the present study was to assess differences in blood-derived DNA methylation patterns (using a high-density array) between individuals with T1DM-ESKD (including individuals requiring dialysis, or those having received a kidney transplant) and individuals with at least 15 years of T1DM and no evidence of kidney disease. Careful and precise phenotyping has provided the ability to minimise differences caused by dialysis treatment; we also evaluated differences in methylation between individuals with T1DM-ESKD who had received a kidney transplant, and thus were receiving immunosuppressive medication, versus those with long duration of T1DM and no evidence of kidney disease.

## Materials and Methods

### Samples

Each participant was recruited as part of the All Ireland-Warren 3-Genetics of Kidneys in Diabetes (GoKinD) United Kingdom Collection. All participants were previously recruited, had White ancestry and provided written informed consent for research. DNA was frozen in multiple aliquots following extraction from whole blood using the salting out method (28) and normalised using PicoGreen quantitation (29) using the CytoFluor® Series 4000 (Applied Biosystems, Thermo Fisher Scientific, CA, USA).

Individuals with both T1DM and ESKD were defined as cases (n=107). These individuals had ≥10 years duration of T1DM alongside a diagnosis of DKD defined as persistent macroalbuminuria (≥500 mg/24hr), estimated glomerular filtration rate (eGFR) <60 mL/min/m^2^ calculated using the Chronic Kidney Disease Epidemiology Collaboration (CKD-EPI) creatinine equation, hypertension (systolic/diastolic blood pressure ≥135/85 mmHg) and ESKD. They were not taking any anti-hypertensive medication. The control individuals had ≥15 years duration of T1DM and no evidence of kidney disease on repeat testing i.e. they all had normal urinary albumin excretion and eGFR >60 mL/min/m^2^ (n=253).

### MethylationEPIC Array

Blood-derived DNA from each participant was accurately quantitated using PicoGreen® prior to normalisation. In total, 800 ng of DNA from each participant was bisulphite treated using the EZ Zymo Methylation Kit (D5002, Zymo Research, CA, USA) following the alternative over-night incubation conditions for use prior to the Illumina® Infinium MethylationEPIC Kit provided in the published protocol. All samples were prepared and analysed using the Infinium MethylationEPIC Kit and BeadChips (Illumina, CA, USA) with no protocol deviations. All samples were processed in a consistent laboratory workstream by the same members of trained staff and methylation arrays were scanned using a dedicated iScan machine with regular monitoring of laser intensity levels. Case and control samples were randomly distributed across the BeadChip arrays. This array is a high-throughput platform which provides quantitative evaluation of methylation levels (β values) with single nucleotide resolution. In total 862,927 sites were examined by the Infinium MethylationEPIC array. The Infinium HD Methylation SNP List was also searched for any SNPs that may have impacted upon the methylation array results (30).

### Quality control

Each resulting .idat file generated from the iScan was assessed using BeadArray Controls Reporter (BACR) Software (Illumina) for quality control (QC). This software assessed the data in connection with a pre-set standard set of controls and evaluated the hybridisation, extension, dye specificity and bisulphite conversion process. An additional QC measure to determine the concordance of average beta values generated for seven duplicate samples was completed using GenomeStudio (Illumina) v1.8, methylation module including a sex check of all included individuals.

Proportional white cell counts (WCCs) were estimated following the Houseman method (31) using the raw .idat files output from the iScan machine. The *minfi* Bioconductor (v3.10) package was utilised to estimate six WCCs, CD8+ T, CD4+ T and CD19+ B lymphocytes, CD56+ natural killer cells, CD14+ monocytes and CD15+ granulocytes using the *estimateCellCounts* function for both the case and control groups, and a Northern Irish general population control group (32).

### Differentially methylated loci analysis

Case and control groups were investigated for dmCpGs using Partek Genomics Suite (PGS) v7.19.1125 (Partek, MO, USA) following functional normalisation. All software was used following the developer’s instructions. Beta values were generated before M values were calculated. DmCpGs were determined using the M values for individuals with T1DM and ESKD compared to controls with long duration of T1DM and no evidence of kidney disease on repeat testing. Parameters were set at false discovery rate (FDR) adjusted p value threshold of ≤×10^−8^ alongside a fold change (FC) ≥±2. Related genes were annotated based on Homo sapiens hg19 genome build using PGS. Four analyses were performed (Table 1), each assessing differential methylation patterns in individuals with T1DM-ESKD to those with T1DM and no evidence of renal disease (Figure 1). Briefly,

1. 107 matched pairs, where both the case and control individuals were matched for age at sample collection (differing by <5 years), sex and duration of diabetes (differing by ≤10 years). Case individuals had a functioning kidney transplant or were receiving dialysis. Controls individuals had T1DM and no evidence of kidney disease.
2. The same 107 individual cases from *Analysis 1* were compared to a larger sample size of (unmatched) control individuals with T1DM and no evidence of kidney disease (n=253).
3. 73 matched pairs for age at sample collection (differing by <5 years), sex and duration of diabetes (differing by ≤10 years). The case subjects were restricted to individuals with a functioning kidney transplant to minimise potential confounding due to differences in medication and renal replacement therapy modalities. These individuals were compared to matched controls with T1DM and no evidence of kidney disease.
4. The same 73 individuals from *Analysis 3* were compared to a larger sample size of (unmatched) control individuals with T1DM and no evidence of kidney disease (n=253).

**Table 1.** Participant group characteristics, per comparison.

|  |  | **Analysis 1\*** | **Analysis 2** | **Analysis 3\*** | **Analysis 4** |
| --- | --- | --- | --- | --- | --- |
| **Cases** | *Number of participants* | 107 | 107 | 73 | 73 |
|  | *Diagnosis* | T1DM with ESKD (transplant recipients or dialysis) | T1DM with ESKD (transplant recipients or dialysis) | T1DM with ESKD (transplant recipients only) | T1DM with ESKD (transplant recipients only) |
|  | *Number of F:M* | 43:64 | 43:64 | 26:47 | 26:47 |
|  | *Average Age (St. Dev.)* | 41.7 years (9.7) | 41.7 years (9.7) | 40.5 years (9.3) | 40.5 years (9.3) |
|  | *Average duration of T1DM (St. Dev.)* | 29.1 years | 29.1 years | 30.1 years | 30.1 years |
| **Controls** | *Number of participants* | 107 | 253 | 73 | 253 |
|  | *Diagnosis* | Long-duration of T1DM with no evidence of kidney disease | Long-duration of T1DM with no evidence of kidney disease | Long-duration of T1DM with no evidence of kidney disease | Long-duration of T1DM with no evidence of kidney disease |
|  | *Number of F:M* | 43:64 | 121:132 | 26:47 | 121:132 |
|  | *Average Age (St. Dev.)* | 41.2 years (10.2) | 41.2 years (9.6) | 40.1 years (9.1) | 41.2 years (9.6) |
|  | *Average duration of T1DM (St. Dev.)* | 26.8 years (7.9) | 26.3 years (7.4) | 25.5 years (7.8) | 26.3 years (7.4) |

\*Analyses 1 and 3: Matched cases and controls for sex, average age ( $\leq 5$ years) and duration of diabetes ( $\leq 10$ years).
Abbreviations: ESKD, end-stage kidney disease; F, females; M, males; St. Dev., standard deviation; T1DM, type 1 diabetes.

**Figure 1:**
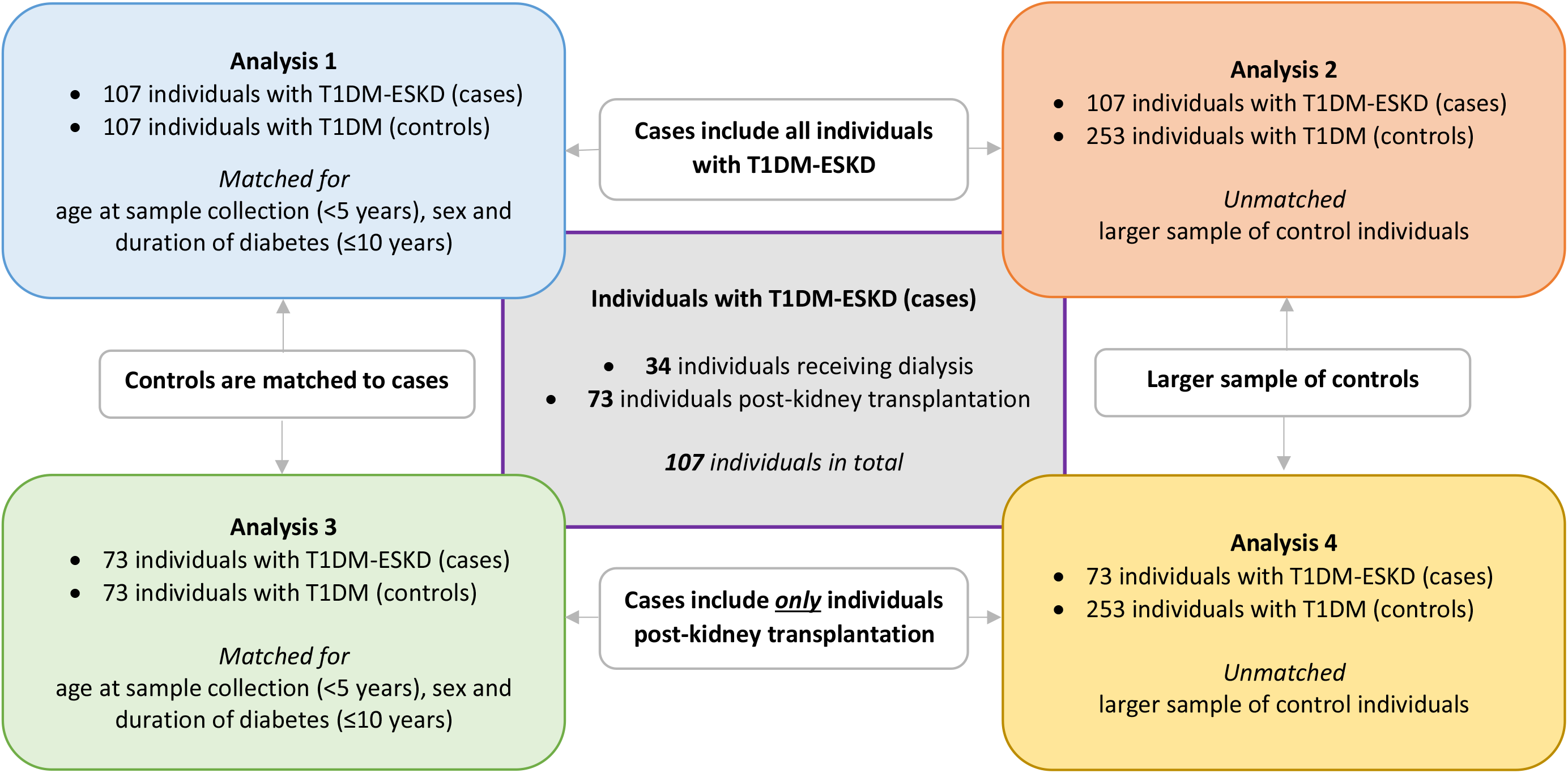
Illustration of groups samples were assigned to in order to complete the analysis.

Resulting dmCpGs which overlapped each of the four analyses were also extracted and displayed in a heatmap which highlighted their degree of hyper- and hypo-methylation.

### Functional gene ontology and network analyses

Gene functionality was examined by gene ontology (GO) and Kyoto Encyclopedia of Genes and Genomes (KEGG) pathway enrichment analysis using PGS. An additional functional analysis was undertaken using STRING v11: protein-protein association networks (33). For each of the four analyses, genes with top-ranked dmCpGs were categorised according to increased or decreased FC.

### Kidney gene expression data

Participants for this study were American Indians with type-2 diabetes (34). Protocol kidney biopsy samples were obtained from this cohort (n = 97). The study was approved by the Institutional Review Board of the National Institute of Diabetes and Digestive and Kidney Diseases and each participant signed an informed consent document. Expression profiling of the kidney biopsies was previously carried out using Affymetrix GeneChips (HumanGenome U133 Array and U133Plus2 Array and Affymetrix Human Gene ST GeneChip 2.1) as reported (35,36), and RNA-seq (Illumina)(13). Expression quantitative trait (eQTL) mapping was performed using an EPACTS (https://genome.sph.umich.edu/wiki/EPACTS) software tool using linear mixed model accounting for hidden familial relatedness, after inverse Gaussian transformation of expression levels, adjusting for age and sex. Significance threshold was set at p≤10^−5^ for eQTL and p<=0.05 for association studies

For morphometry measures, kidney biopsy tissue was prepared for light and electron microscopy studies according to standard procedures (37–39). The following glomerular structural parameters were measured by unbiased morphometry on electron microscopy images as described elsewhere (37,38,40): glomerular basement membrane (GBM) width (41,42), mesangial fractional volume (VVMES) (41,42), glomerular filtration surface area per glomerular volume (SV) (41,42), percentage of endothelial fenestrations (P_FEN) (43) and the fractional podocyte volume per glomerulus (VVPC) (44). Cortical interstitial fractional volume (VVINT) (39) were estimated using light microscopy with all specimens embedded in paraffin.

### Replication cohorts for the methylation data

Replication cohort was comprised of 327 peripheral blood samples obtained from Pima Indians (24). Blood samples for DNA isolation and cytosine methylation analysis were collected at the baseline examination from subjects. EGFR measures were available from clinical records before and after the nephrectomy. Among these subjects, 318 had longitudinal eGFR measurements with mean follow-up time of 9.8 years with a standard deviation (SD) of 3.9 years. The best linear unbiased predictor (BLUP) model was used to calculate eGFR slope for each subject.

Human kidney samples (n=91) were obtained from surgical full/partial nephrectomies. Kidney samples were ~0.5 cm in diameter and were surrounded by at least 2 cm of normal tissue margins. Clinical data was obtained at the time of the sample collection and for a subset of samples eGFR measures were available from clinical records before and after the nephrectomy (*n* = 69). Only subjects with longitudinal eGFR measurements for at least three months post nephrectomy were included for the analysis. The mean timespan of follow-up was 2.4 years (SD = 1.5 years).

DNA methylation was detected using Illumina 450K arrays and values were extracted using the minfi package. Several quality control measures were carried out and after the removal of CpG probes that (1) were in the proximity of regions with genetic variations, (2) were located on the sex chromosomes, (3) were known to cross-hybridize to other locations, (4) had poor detection p values (p > 0.01) or (5) were control probes, 321,473 CpG probes remained in the blood replication cohort. The same pre-processing was performed for the kidney replication cohort which resulted in 355,141 CpG probes remaining.

In blood replication cohort, linear regression models were used to determine the association between DNA methylation, eGFR and eGFR slope, with covariates including sex, age, duration, mean arterial blood pressure, HbA1c, batch, converse and estimated cell fraction. Association between cytosine methylation level, eGFR and interstitial fibrosis was determined by linear regression models adjusted by age, sex, race, diabetes, hypertension, batch effect, bisulphite conversion efficiency and degree of lymphocytic infiltrate on histology. A p value of <0.05 was used to determine significance in this replication cohort.

## Results

The Illumina Infinium MethylationEPIC BeadChip array examines 862,927 CpG sites. Each resulting .idat file generated from the iScan was assessed using Illumina’s BACR software. This software assessed the data in connection with a pre-set standard set of thresholds (45) which are indicated alongside the BACR data for each analysis. QC was completed with no significant difference in intensity levels detected. On average, fewer than 30 probes failed per sample. Houseman estimates (31) were calculated for the proportional WCCs of each sample and the results were included as a Supplementary Table (ST) for each of the four analyses; ST2, ST8, ST15 and ST21. Proportional WCCs for each group were compared with a general population control group, NICOLA (Northern Ireland Cohort for the Longitudinal Study of Ageing).

Blood-derived DNA from 360 individuals was included in this analysis. Of these participants, 253 were control individuals with a long duration of T1DM and no evidence of kidney disease upon repeated testing. The mean age of these participants was 41 years and with an average duration of diabetes of 26 years and without anti-hypertensive medication usage. The remaining 107 individuals were diagnosed with T1DM-ESKD, 73 of whom had received a kidney transplant. The average age of these participants was 41 years. The participant characteristics are provided in Table 1.

This study was conducted using four complementary analyses; matching of the participants was required for *Analyses 1* and *3*. Individuals were matched for age at sample collection (differing by <5 years), sex and duration of diabetes (differing by ≤10 years). Concordance plots were drawn for seven duplicate samples; average r^2^= 0.99; Supplementary Figure (SF) 1.

### Analysis 1: Individuals with T1DM and ESKD compared to <u>matched</u> control individuals with T1DM and no evidence of kidney disease

This first analysis included 107 matched pairs, where both the case and control individuals were matched on sex and age at diagnosis (Table 1). The BACR QC report is included within ST1. Proportional WCCs were estimated for all included participant blood samples. ST2 shows the average WCCs for *Analysis 1* and the results of a t-test comparing these between the two sample groups. Two cells, granulocytes and monocytes showed significant differences at the 0.05 level, the most significant of which was 0.04; but after correction for multiple testing of the WCC analyses, these results are not significant (p>0.002).

Comparison of the methylation patterns between cases and controls identified 4,391 top-ranked dmCpGs (FDR p≤×10^−8^, ST3). Two genes, *ARID5B and SEPTIN9*, contained ≥10 top-ranked dmCpGs. When the stringency was increased to include a larger change in methylation between cases and controls (FDR p≤×10^−8^ and FC ±2), 490 CpG sites remained significant (ST4) of which 332 were gene centric. Of these, 22 genes contained at least two dmCpGs including *HDAC4*, *ITGAL*, *LY9*, *PBX1*, *RPTOR*, *RUNX3* and *SEPTIN9* (ST4).

To further assess the functional significance of these changes in DNA methylation between case and control groups, a GO enrichment analysis was undertaken. This analysis assessed the biological processes, cellular components and molecular functions of the genes within which the 490 top-ranked CpG sites were located (FDR p≤×10^−8^ and FC ≥±2). A total of 325 GO functions had an enrichment score ≥4 (p<0.01, ST5 and SF2). The processes with the top enrichment scores include cell surface receptor signalling pathway, immune system process and positive regulation of immune system processes, lymphocyte activation and T cell activation. The KEGG pathway database was searched to identify key pathways linked to the genes where the top-ranked dmCpGs were located. Eleven pathways were identified with an enrichment score of ≥2 and p≤0.01 (ST6). This analysis of differentially methylated genes returned pathways including Th17 cell differentiation, human T-cell leukaemia virus 1 infection, and natural killer cell mediated cytotoxicity.

### Analysis 2: Individuals with T1DM and ESKD compared to a larger cohort of <u>unmatched</u> control individuals with T1DM and no evidence of kidney disease

This second analysis included the same 107 individual cases from *Analysis 1* with a larger sample size of (unmatched) control individuals with T1DM and no evidence of kidney disease (Table 1). The BACR QC report and proportional WCCs for this analysis are included within ST7 and ST8 respectively.

Following the same analysis path as previously described, comparison of methylation between case individuals and control individuals identified 13,983 top-ranked dmCpGs (FDR p≤×10^−8^; ST9). Two genes *ETS1* and *UBAC2* contained over 20 dmCpGs. When the stringency levels were increased (FDR p≤×10^−8^ and FC ±2), 1,112 CpG sites remained, of which 768 were gene centric (ST10). *SEPTIN9* contained the largest number of dmCpGs (n=5) with the criteria set to include FC±2. Comparison of results from *Analyses 1* and *2* (FDR p≤×10^−8^ and FC ±2) identified 325 dmCpGs (223 within genes) which overlapped. Each dmCpG demonstrated the same direction of effect (ST11).

GO enrichment analysis was similarly undertaken to assess the functional significance of the 1,112 significant DNA methylation alterations between the case and control groups. A total of 505 GO functions had an enrichment score ≥4, alongside p<0.01 (ST12 and SF3). The processes with the top enrichment scores include several linked to immune responses including regulation of immune system processes and lymphocyte activation. The KEGG pathway analysis revealed 16 pathways (enrichment score of ≥2, and p≤0.01; ST13), including cancer, acute myeloid leukaemia and natural killer cell mediated cytotoxicity.

### Analysis 3: Individuals with T1DM and ESKD who have received a kidney transplant compared to <u>matched</u> control individuals with T1DM and no evidence of kidney disease

The inclusion criterion for the case subjects was restricted to individuals with a functioning kidney transplant for the third analysis. Only blood-derived DNA samples from individuals who had received a kidney transplant were included to minimise potential confounding due to differences in medication and renal replacement therapy modalities (n=73). The methylation status of the CpG sites for these individuals was compared to matched controls with T1DM and no evidence of kidney disease (n=73). The BACR QC report and proportional WCCs for this analysis are included within ST14 and ST15 respectively.

In total, 1,518 top-ranked dmCpGs were different between cases and controls (FDR p≤×10^−8^; ST16). Of those, 132 of the top-ranked dmCpGs remained when the stringency levels were increased (FDR p≤×10^−8^ and FC ±2), 81 of which were gene centric including two within *MTURN* and *UPF3A (ST17)*. Additional GO enrichment and pathway analyses were undertaken for the 132 top-ranked genes in which the dmCpGs were located (FDR p≤×10^−8^ and FC±2) to assess their functional significance. In total, 75 GO functions were enriched with a score ≥4, alongside p<0.01 (ST18, SF4). The KEGG pathway analysis showed one result - the TGF-β signalling pathway with an enrichment score of ≥2 and p≤0.01, ST19).

### Analysis 4: Individuals with T1DM who had received a kidney transplant compared to <u>unmatched</u> control individuals with T1DM and no evidence of kidney disease

The methylation status of the CpG sites for individuals who had received a kidney transplant was compared to unmatched controls with T1DM and no evidence of kidney disease (n=253). The BACR and proportional WCCs QC report are included within ST20 and ST21 respectively. In total, 13,739 top-ranked dmCpGs were identified between both groups (FDR p≤×10^−8^, ST22). Of these, 723 dmCpGs were gene centric. When the stringency levels were increased (FDR p≤×10^−8^ and FC ±2), 1,082 CpG sites remained of which 723 were located within genes (ST23). Seven genes including *ACAD8*, *LIME1*, *RPTOR* and *SEPTIN9* each contained three dmCpGs. In total, 78 dmCpGs (48 were within genes) overlapped between *Analyses 3* and *4* (FDR p≤×10^−8^ and FC ±2, ST24).

GO enrichment and pathway analyses were completed for the genes in which the 1,082 top-ranked dmCpGs were located (FDR p≤×10^−8^ and FC ±2). A total of 679 GO functions had an enrichment score ≥4 and p<0.01 (ST25 and SF5). The processes with the top enrichment scores include regulation of cell activation, enzyme binding and immune system processes. Fourteen KEGG pathways were identified (enrichment score of ≥2, and p≤0.01; ST26) including cancer and platelet activation.

### Significantly associated dmCpGs from Analyses 1-4

Thirty-six dmCpGs were identified as significantly different between cases and controls in each of the four analyses (FDR p≤×10^−8^ and FC ±2; ST27 and SF6). The direction of fold change was consistent for each *Analysis*.

### Search for overlapping SNPs

Previous studies have reported that SNPs could potentially impact on methylation status (46). To assess this, we searched the Infinium HD Methylation SNP List (30) for any SNPs that could potentially impact the methylation array results if present in the test population. Of the top-ranked dmCpGs from this analysis (FDR p≤×10^−8^, FC ±2), five single CpG sites have the potential to be affected by SNPs (ST28). All but two of these SNPs, rs4788986 (*SEPTIN9*) and rs742232 (*RUNX3*) are very rare in European populations and would therefore be unlikely to impact in this study. Of note, multiple dmCpGs were identified in both *SEPTIN9* and *RUNX3* genes with functional support provided for both these genes, making it improbable that these genes are falsely identified.

### STRING Functional analyses

Functional network analyses were undertaken using STRING v11 (33) for the list of genes in which the top-ranked dmCpGs were located. Those which showed an increase in FC in the individuals with T1DM-ESKD compared to individuals with T1DM were analysed separately to those genes which showed a decrease in FC. All pathway interactions are shown in SFs7–14.

### Functional data results

The genes in which top-ranked dmCpGs were located, which also had the most biological plausibility for ESKD, were assessed for eQTLs in an American Indian population with T2DM(47). EQTLs demonstrated support for eight of the top-ranked dmCpG genes; *CUX1*, *ELMO1*, *HDAC4*, *PIM1*, *PRKAG2*, *PTPRN2*, *RUNX3*, *SEPTIN9* (p≤10^−5^, ST29). Functional support was also sought from existing glomerular and tubular expression data alongside eGFR and fibrosis in both blood and kidney biopsy tissue using data ascertained from the 450K array (all p<0.05)(24,48). Seven differentially methylated genes were replicated in the glomerular database and six in the tubular (ST29; Table 2).

**Table 2.**
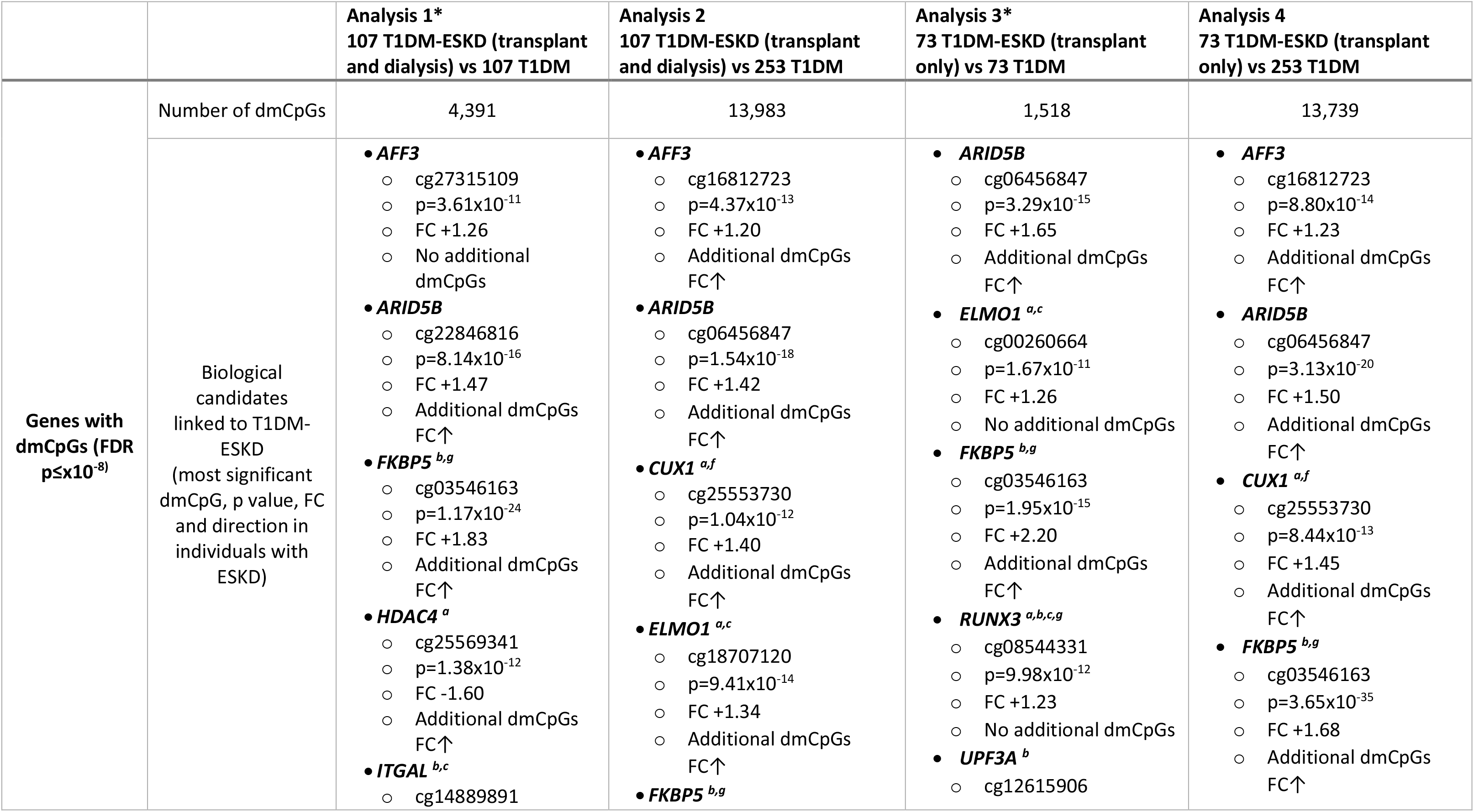
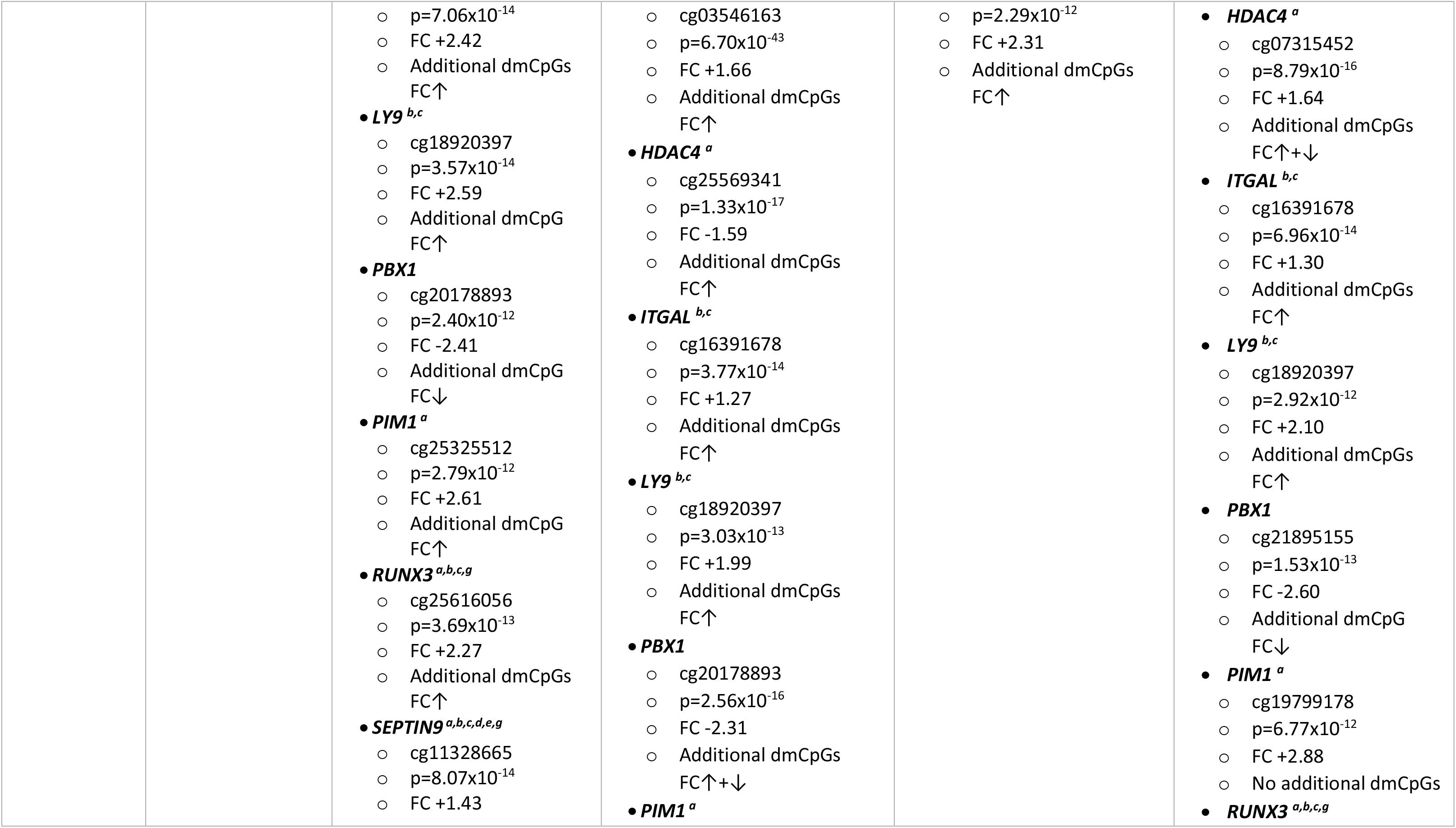
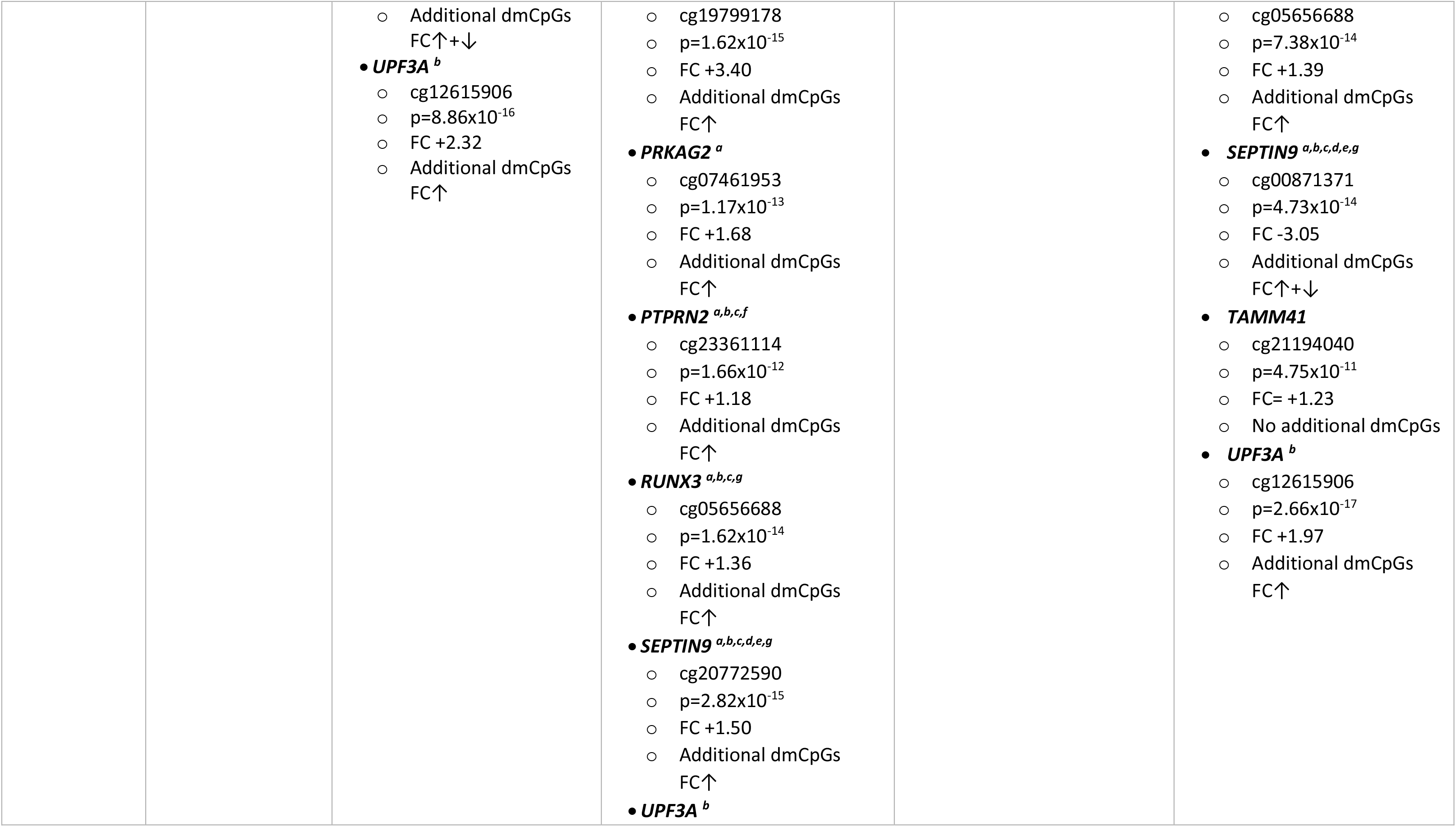
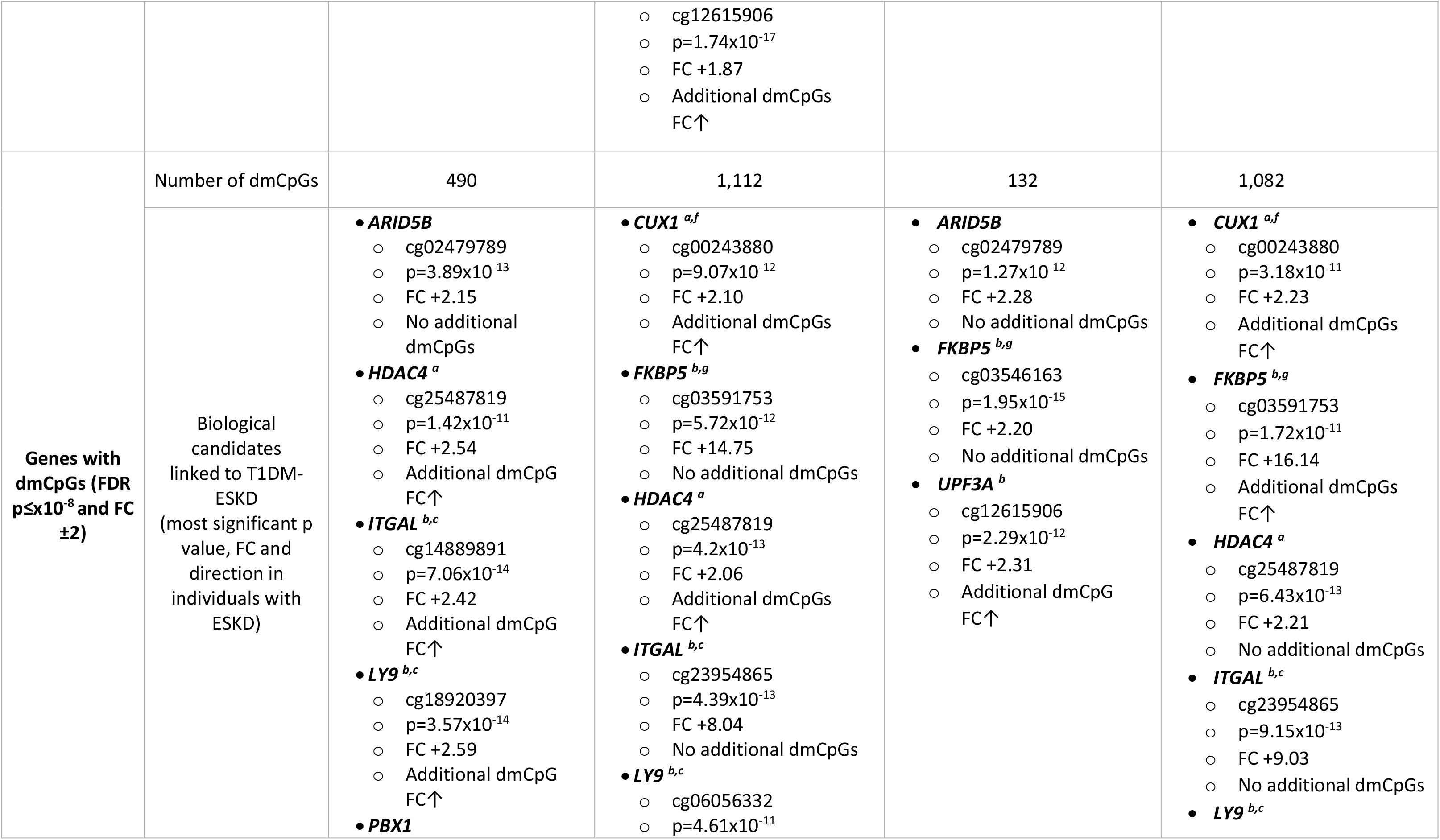
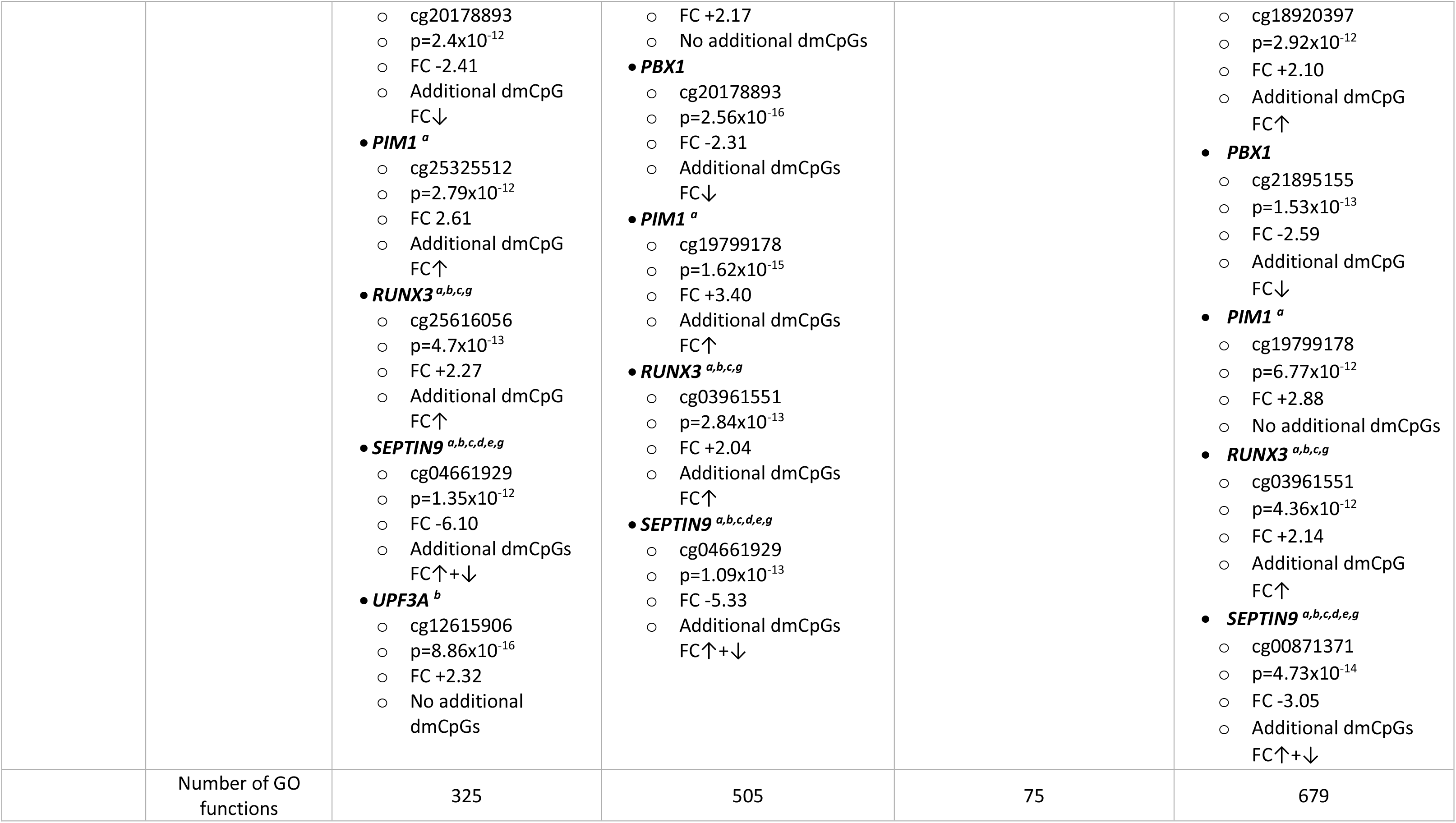
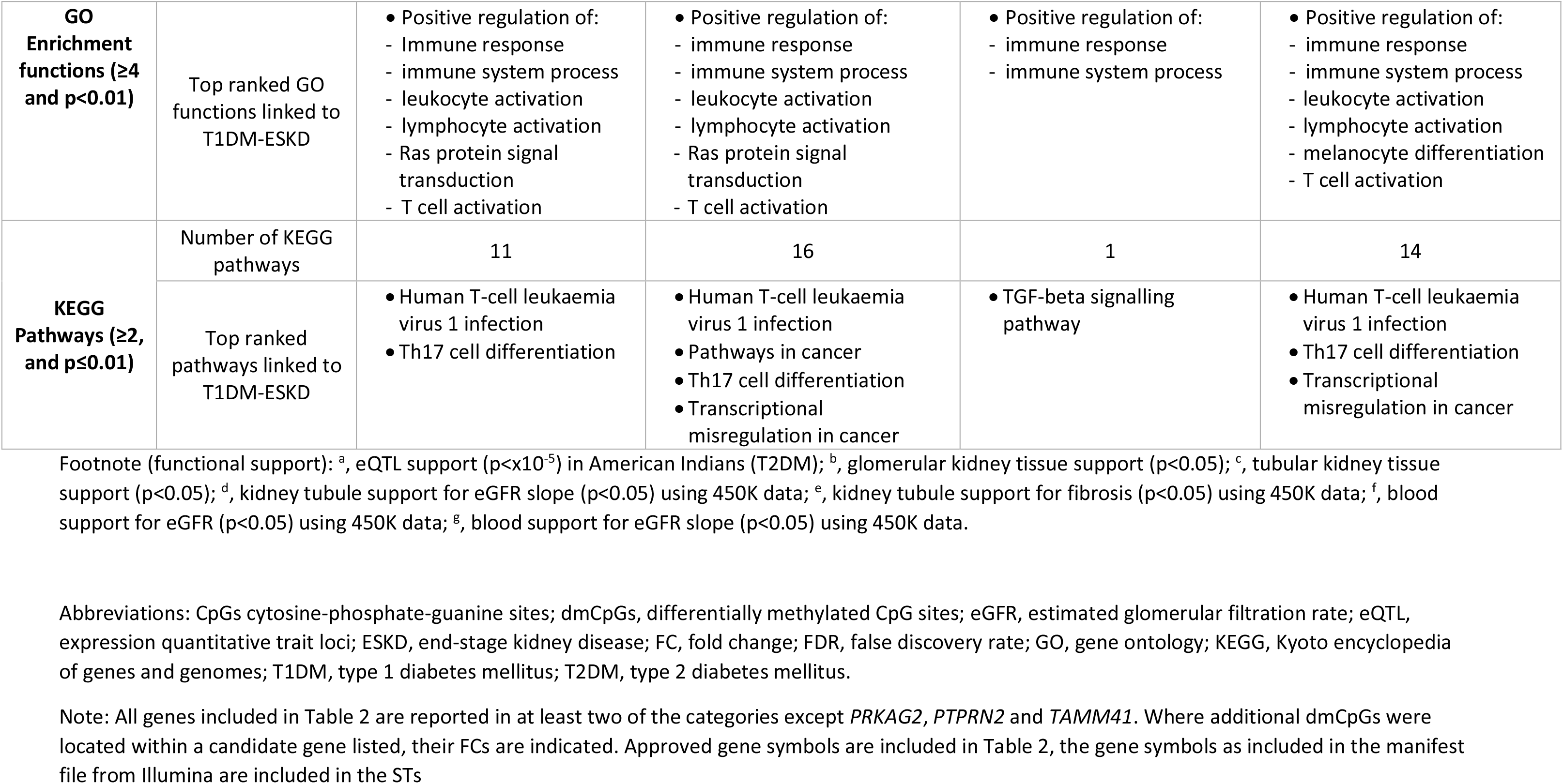
Summary results, highlighting top-ranked dmCpGs, GO enrichment functions and pathways per analysis.

### Replication analysis

We have examined the association between the methylation of top ranked CpGs and kidney function from this study and previous studies which had been performed on subjects with T2DM, including cohorts with blood sample methylome analysis and others where methylation was studied in micro-dissected human kidney tubule samples. We confirmed nominally significant association with kidney function at the *CUX1*, *PTPRN2* and *SEPTIN9* locus and with kidney function decline at *AFF3*, *HDAC4* and *SEPTIN9* regions. Furthermore, the methylation level at the *AFF3*, *CUX1* and *PIM1* regions correlated with the degree of fibrosis in micro-dissected human kidney tissue samples.

### Summary

A summary of results is included within Table 2, SF15 and SF16, including specific details regarding the strongest biologically plausible candidates linked to T1DM-ESKD, the GO functions and pathways. Table 2 highlights the top-ranked genes and pathways of interest and states the number of significant results from each section of the analysis.

## Discussion

Epigenetic alterations provide a dynamic link between genetic background and environmental exposures. These alterations have been proposed to play an important role in kidney disease (20,23) and T1DM (49). Previous research assessing T1DM-DKD (19–22,50) has identified dmCpGs using bisulphite pyrosequencing and either the Illumina Infinium 450K or 27K methylation arrays. This manuscript describes differences in DNA methylation patterns between individuals with T1DM and those with T1DM-ESKD, utilising the MethylationEPIC array technology.

Several top-ranked dmCpGs identified through this study showed an increase in FC related to the more severe phenotype of T1DM-ESKD compared to the T1DM control population.

The genes from each analysis with the most biological plausibility having assessed the literature, are included in Table 2 and SFs 14 and 15. Of the 16 genes in which dmCpGs were located with a significance level of FDR p≤×10^−8^, (SF14), 11 retained this status when the stringency level was increased (FDR p≤×10^−8^ and FC±2; SF15). DmCpGs were located within *FKBP5* and *RUNX3* in each of the analyses where FDR p≤×10^−8^. When the stringency was increased (FDR p≤×10^−8^ and FC±2), they remained top-ranked genes in all but *Analyses 1* and *3* respectively.

*FKBP5*, also known as FK506 binding protein 51, acts as a co-chaperone of hsp90 to aid the modulation of glucocorticoid receptor sensitivity in response to stress (51,52). Polymorphisms in this gene lead to an extended stress hormone response following exposure (51). Recently, genome-wide analyses of human blood found associations between *FKBP5* mRNA and a pro-inflammatory profile (53). Moreover, aberrant *FKBP5* methylation has previously been implicated in the pathology of numerous diseases, particularly in diseases common in older populations. This hypermethylation is exemplified in myocardial infarction (53) and conditions such as T2DM (54) and CKD (20). All top-ranked dmCpGs within this gene were located in either the gene body or 5’ UTR and showed increased methylation for individuals with T1DM-ESKD. As a hallmark feature of CKD is persistent, low to moderate levels of circulating inflammatory markers (55), with distinguishing features such as nephron loss with subsequent acceleration of organ fibrosis, further study is required to determine if *FKBP5* plays a mechanistic role in CKD development.

The runt related transcription factor 3 (*RUNX3*) plays a downstream role in the TGF-β signalling pathway (56). Its suppression has been implicated in tumour growth, migration and invasion (57). In 2019, Cen et al., reported that higher methylation levels in *RUNX3* were associated with a shorter renal cell carcinoma survival time (58). Additionally, they suggested independent predictors of heightened methylation levels of this gene included the presence of intra-tumour vascularity. We identified increased FC in dmCpGs within the *RUNX3* gene body and the UTRs in association with T1DKD-ESKD.

As indicated in SF15, a further six genes, *HDAC4*, *ITGAL*, *LY9*, *PBX1, PIM1* and *SEPTIN9* had top-ranked dmCpGs from three of the four analyses. These genes did not reach statistical significance in *Analysis 3*, which had the smallest population which compared DNA methylation patterns in 73 individuals with T1DM-ESKD (transplant only) to 73 individuals, matched, with T1DM.

Histone deacetylases (HDACs) are a group of enzymes that are characterised into three defined classes, known as I, II and III (59,60). They each have roles in removal of acetyl groups from histone and non-histone proteins, chromatin condensation and transcriptional repression (59,60). They have the ability to impact cellular function via both epigenetic and non-epigenetic mechanisms (60). *HDAC4* (a member of class II) was reported to reduce kidney injury during *in vivo* animal studies (59–61). Additionally, *HDAC4* was found to be over-expressed in kidney epithelial cells of a murine kidney fibrosis model (62). Subsequent treatment with HDAC inhibitors demonstrated that the development and progression of kidney fibrosis can be inhibited by suppressing the activation and expression of numerous pro-fibrotic molecules, such as fibronectin and collagen 1 (62). In the present study, *HDAC4* showed an increased FC for all dmCpGs except cg25569341 in individuals with ESKD undergoing dialysis or who had undergone a transplant.

Integrins are integral membrane proteins which are heterodimeric in nature. They are comprised of alpha and beta chains which together form the integrin lymphocyte function-associated antigen-1, expressed in leukocytes. *ITGAL* encodes the alpha L chain and its expression has been previously linked to renal cell carcinoma (63). A second report has demonstrated that DNA methylation of the *ITGAL* gene is heavily methylated in fibroblasts and demethylated in T lymphocytes (64). The top ranked dmCpGs present within the gene body or north shelf of *ITGAL* showed an increase in FC in individuals with T1DM-ESKD who were immunosuppressed following transplantation, compared to those with T1DM with no evidence of kidney disease.

Lymphocyte antigen 9 *(LY9)* is a member of the signalling lymphocyte activation molecule family receptor and is involved in immune responses. In 2019, Parikova et al. (65) reported expression of *LY9* to be significantly increased in individuals receiving long-term dialysis compared to those who had received dialysis over a shorter time period. Our findings support these observations showing an increased FC in *LY9* dmCpGs located within either the gene body or 5’ UTR in individuals with T1DM-ESKD compared to those with no kidney disease.

*PBX1* encodes a nuclear protein within the PBX homeobox family of transcription factors. Through this it can affect the expression of several genes including those which regulate insulin action and glucose metabolism (66). In the current study, dmCpGs within the *PBX1* gene body showed a consistent decrease in FC at the highest stringency level (FDR p≤×10-8, FC ±2) in individuals with T1DM-ESKD across each of the four analyses. Previous analyses of this gene have linked differential methylation patterns to higher birth weight-for-gestational age (67) and a translocational rearrangement of this gene and *TCF3/E2A* has been associated with B-cell acute lymphoblastic leukaemia (68). Additionally, *PBX1* haploinsufficiency has been linked to congenital anomalies of the kidney and urinary tract (69), reported to be involved in the proliferation of cells in renal cell carcinoma (70) and has been recognised as a candidate gene for T2DM (66,71).

*PIM1* belongs to the serine/threonine kinase family (72) and its overexpression has previously been implicated in diseases such as ovarian cancer (73) and breast cancer (74). Overexpression of *PIM1* appears to influence cancer development in three ways; preventing apoptosis, enhancing cellular proliferation and through promoting genomic instability (75). More specific to kidney research, *PIM1* is aberrantly overexpressed in renal cell carcinoma (76) and lupus nephritis (77). In this analysis we identified top-ranked dmCpGs present within the 3’ UTR and north shore of *PIM1* which resulted in an increase in the FC in individuals with T1D-ESKD.

Septins are a group of GTPase proteins that play a role in cytoskeleton organisation through their links with microtubules and actin filaments (78). Although some septins are associated with the advancement of kidney fibrosis (78), *SEPTIN9* specifically has been implicated in a host of disease pathologies, such as prostate cancer (79), colorectal cancer (80), liver fibrosis (81) and T2DM (82). Moreover, *SEPTIN9* overexpression has been shown to promote kidney epithelial cell migration (83). Additionally Dayeh et al. reported *SEPTIN9*, alongside *PTPRN2,* as one of the top-ranked differentially methylated genes when comparing pancreatic islets from individuals with and without T2DM (84). In this analysis, 20 dmCpGs were located in *SEPTIN9,* 17 of which displayed an increase in FC in individuals with T1DM-ESKD.

An additional three top-ranked genes which contained dmCpGs were common in two analyses; *ARID5B* and *UPF3A*, which were identified from *Analyses 1* and *3*, and *CUX1* which resulted from *Analyses 2* and *4* (FDR p≤×10^−8^ and FC±2; SF15).

Each of the dmCpGs within *ARID5B* were located within the gene body and all were consistently increased in cases with ESKD (both chronic dialysis and transplant recipients) compared with controls with long duration of T1DM and no evidence of kidney disease. It has been previously demonstrated that *ARID5B* has a regulatory role in the phenotypic change of smooth muscle cells and SNPs within *ARID5B* have been linked to T2DM and coronary artery disease (85). Differential methylation of a six-probe region spanning 99 base pairs within *ARID5B* gene has also been reported in Alzheimer’s disease (86). *ARID5B* is also known to contribute to cell growth, the differentiation of B-lymphocyte progenitors and has additionally been linked to acute lymphoblastic leukaemia (87–89).

*UPF3A* encodes a protein involved in mRNA nuclear export and mRNA surveillance, it has a crucial role in downregulating aberrant mRNAs (90). Gotoh et al. in 2014 reported that this gene alongside 14 others had significantly reduced mRNA expression levels in renal cell carcinoma compared to non-cancerous kidney cortex tissue (91). Top-ranked dmCpGs present within *UPF3A* consistently displayed an increase in FC within individuals with T1DM and ESKD.

*CUX1* is a protein coding member of the homeodomain family of DNA binding proteins and is involved in cell cycle regulation and kidney development through the inhibition of p27 which promotes cell proliferation in the nephrogenic zone (20,92,93). Previously, significant DNA methylation alterations have been reported in CKD where more than one dmCpG site was located within *CUX1* compared to individuals with no evidence of kidney disease (20). Additionally, genetic abnormalities within *CUX1* have been linked to polycystic kidney disease (94) and myelodysplastic syndrome (95) in mouse models. Significant dmCpGs from *Analyses 2* and *4* were located within the body of this gene which additionally showed an increase in FC in individuals with T1DM-ESKD, but this gene has not previously been explored in individuals post kidney transplant.

Lastly, an additional five genes with biological plausibility for ESKD had statistical significance for this phenotype but did not show a large change in methylation (FC±2); *AFF3*, *ELMO1*, *PRKAG2*, *PTPRN2* and *TAMM41* (SF14, FDR p≤×10^−8^). These genes contained dmCpGs from either one (*PRKAG2*, *PTPRN2* and *TAMM41*), two (*ELMO1*) or three analyses (*AFF3*).

In 2012, the GENIE consortium conducted a meta-analysis of GWAS in T1D-DKD, which revealed an intronic SNP (rs7583877) located in the AF4/FMR2 family member 3 (*AFF3*) gene as significantly associated with ESKD (14,96). Functional studies have indicated that *AFF3* influences kidney tubule fibrosis through the TGF-β1 pathway (14). The findings from our study show increased methylation levels in the dmCpGs within the body of *AFF3* in the individuals with T1D-ESKD, which could result in decreased gene expression. A link between DNA methylation of this gene and T1DM has previously been considered as a mediator of the genetic risk (97). Each of the top-ranked dmCpGs identified from this analysis were present within the body of *AFF3* with increased methylation in individuals with T1DM-ESKD.

*ELMO1* encodes a member of the engulfment and cell motility protein family and has been previously linked to T2DM (98–100), hepatocellular carcinoma (101) and inflammatory arthritis (102). Previous methylation analyses of this gene have shown associations with gastric cancer (103) and CKD (20). The eight top-ranked dmCpGs within *ELMO1* were present within various regions of the gene and each dmCpG site reported an increase in the FC in individuals with T1DM-ESKD. One hypermethylated dmCpG site, cg01119452, had been previously reported in 2014, where it also showed hypermethylation in its association with CKD (20).

*PRKAG2* is an important regulator of cellular energy status and has previously been associated with eGFR (104). SNP rs7805747 located in *PRKAG2* has been reported in association with both CKD and serum creatinine at genome-wide significance level (105). Differential methylation within this gene has also previously been reported in association with CKD (20). The top-ranked dmCpGs were present within the 5’ UTR or body of *PRKAG2* and each showed an increase in FC in individuals with T1DM-ESKD compared to those with no evidence of kidney disease.

Identified as an auto-antigen in diabetes, *PTPRN2* has previously been linked to CKD (20,106), fasting plasma glucose and obesity (107). *PTPRN2* encodes islet antigen (IA)-2β and together with IA-2, these are integral membrane proteins of dense core vesicles which are expressed throughout the body in neuroendocrine cells (108). *PTPRN2* was reported by Dayeh at el. as second of the top-ranked differentially methylated genes in a comparison of pancreatic islets from individuals with and without T2DM (84). Each of the dmCpGs present within *PTPRN2* showed a consistent increase in FC in association with T1DM-ESKD.

The function of *TAMM41* in higher vertebrates still remains largely undetermined, yet it is known to play a critical role in yeast cell mitochondrial membrane maintenance (109). Using zebrafish models of human CVD, it was determined that the developing heart overexpressed *tamm41* (109,110). Furthermore, CRISPR/Cas9 mediated knockout of the t*amm41* gene resulted in immature heart valve formation (109). Differential methylation in *TAMM41* has previously been reported in both DKD and ESKD (22).

Several of the top-ranked pathways and genes defined by dmCpGs have been previously linked to leukaemia. This is not unexpected as an elevated risk of leukaemia during dialysis and after transplant failure has previously been reported in an Australian and New Zealand population (111,112). Although rare, Alfano et al. suggested that leukaemia can occur during the post-transplant period (113).

In the absence of a complementary replication cohort with available data, further support for top-ranked genes was sought from eQTL analysis from an American Indian cohort where the individuals had T2DM and known renal status (47). eQTL analysis supports associations for eight of the top-ranked genes; *CUX1*, *ELMO1*, *HDAC4*, *PIM1*, *PRKAG2*, *PTPRN2*, *RUNX3*, *SEPTIN9* (p≤10^−5^, ST29). Functional support was also generated for these top-ranked markers in kidney tissue (glomerular tissue and tubule tissue from two cohorts (47,48)) alongside replication in additional blood derived DNA from individuals with kidney function assessed using the 450K array (unpublished data)(48). Unfortunately, the 450K array for DNA methylation includes <25% of the top-ranked dmCpGs identified by our study which used the more comprehensive EPIC array. It is not unexpected that the dmCpGs did not correlate across all top-ranked results due to the differences in material and technologies employed.

Through this analysis we have demonstrated that similar association results are reported for individuals who have received a kidney transplant or persons who are receiving dialysis. This clearly shows that transplant recipients can be analysed alongside individuals receiving dialysis to increase the power of future EWAS for ESKD. The results; genes, p-values and FC directions are consistent across each of the four analyses. *RPTOR* is the only top-ranked gene linked to immunosuppressant medication for transplants (114).

We have reported associations with several biologically plausible genes. Employing a considered approach, including four analyses (two of which were matched and two were unmatched), we show broad overlap in results from the analyses. This indicates that for future large-scale studies, it is not essential to stratify this analysis by age and sex matching the participants, which will facilitate a larger, more powerful case-control design.

### Strengths and Limitations

Overall, this study has several strengths. We have carefully defined phenotypes for each analysis, ensuring individuals were well matched for paired *Analyses 1* and *3*, with a larger cohort of individuals employed for unmatched *Analyses 2* and *4. Analyses 3* and *4* represented an extension of the initial analyses to evaluate the differences between individuals with T1DM and DKD compared to those with a more extreme phenotype who had progressed to T1DM-ESKD and had received a kidney transplant.

We have extended previous investigations through assessing T1DM-ESKD using the most cost-effective high-density methylation array available, the Infinium MethylationEPIC (115). Methylation signatures were assessed using peripheral blood samples, while considering estimates of proportional WCCs and cell heterogeneity. Furthermore, only top-ranked dmCpGs that met a significance level of FDR p≤×10^−8^ were reported, a threshold previously reported to reduce the rate of false-positives in studies which use the Infinium MethylationEPIC array (116). This array solely focuses on the presence of methylation at CpG sites and does not take non-CpG site methylation into consideration.

This study is limited in not having a complementary replication cohort, however we took the approach that a larger discovery cohort using the most comprehensive array commercially available was the optimal study design given similar kidney transplant cohorts do not have EPIC array data available for replication. While we have provided details of all dmCpGs with FDR p≤×10^−8^ and FC±2 in supplementary material, this manuscript focuses mainly on genes that have previously been linked to similar phenotypes in the literature. A larger-scale multi-omic analysis incorporating genetic variation, epigenetic alterations and gene expression would be required to further determine the markers of interest for this phenotype and improve understanding of the biological mechanisms involved. Additional investigations could be undertaken to assess this phenotype in different ethnicities, sex-specific effects, and analyses could be conducted to compare these results with those derived from kidney biopsy samples.

### Conclusion

Epigenetic alterations, unlike genetic changes, are potentially reversible, offering opportunistic therapeutic interventions. We have reported associations between dmCpGs and genes with T1DM-ESKD, several of which suggest complementary genetic and epigenetic influences to alter gene expression. Eight top-ranked genes also showed eQTL support in a T2DM American Indian cohort and 13 were supported by gene expression and / or methylation data from kidney tubule or glomerular tissues. Additional prospective studies may help identify whether the underlying methylation influences are causal or consequential.

The identification of unique epigenetic profiles associated with developing ESKD could highlight additional biological mechanisms to study kidney disease. Epigenetic profiles may also help to identify patients at greater risk for progression to ESKD. Targeting healthcare resources to individuals at highest risk for ESKD remains an important clinical goal.

## List of abbreviations

BACR: Bead Array Controls Reporter
CKD: chronic kidney disease
DKD: diabetic kidney disease
dmCpGs: differentially methylated
eGFR: estimated glomerular filtration rate
eQTL: expression quantitative trait loci
ESKD: end-stage kidney disease
EWAS: epigenome-wide association study
FC: fold change
FDR: false discovery rate
GBM: glomerular basement membrane width
GENIE: GEnetics of Nephropathy an International Effort
GO: gene ontology
GoKinD: Genetics of Kidneys in Diabetes
GWAS: genome wide association studies
HbA1c: haemoglobin A1c
HDACs: histone deacetylases
KEGG: Kyoto Encyclopedia of Genes and Genomes
NICOLA: Northern Ireland Cohort for the Longitudinal Study of Ageing
P_FEN: percent of endothelial fenestration falling on the peripheral glomerular basement membrane
PGS: Partek Genomics Suite
QC: quality control
SF: supplementary figure
ST: supplementary table
SV: surface volume of peripheral glomerular basement membrane per glomerulus
T1DM: type 1 diabetes mellitus
T2DM: type 2 diabetes mellitus
VVINT: cortical interstitial fractional volume
VVMES: mesangial fractional volume
VVPC: volume fraction of podocyte cell per glomerulus
WCCs: white cell counts

## Acknowledgements

We thank all individuals who participated in this study and acknowledge all physicians and nurses who were involved in their recruitment.

The GENIE_UK data includes samples recruited as part of the Warren3/U.K. GoKinD Study Group, jointly funded by Diabetes UK and the Juvenile Diabetes Research Foundation and includes the following individuals: Professor A.P. Maxwell, Dr. A.J. McKnight, Dr. D.A. Savage (Belfast); Dr. J. Walker (Edinburgh); Dr. S. Thomas, Professor G.C. Viberti (London); Professor A.J.M. Boulton (Manchester); Professor S. Marshall (Newcastle); Professor A.G. Demaine and Dr. B.A. Millward (Plymouth); and Professor S.C. Bain (Swansea).

## Funding Details

LJS is the recipient of a Northern Ireland Kidney Research Fund Fellowship (NIKRF).

The GEnetics of Nephropathy an International Effort (GENIE) through the Medical Research Council (MC_PC_15025), Public Health Agency R&D Division (STL/4760/13), Science Foundation Ireland (SFI15/US/B3130), NIH R01_DK081923 and R01_DK105154. LJS, KA, APM and AJM, are also supported by Science Foundation Ireland and the Department for the Economy, Northern Ireland US partnership award 15/IA/3152. The data referenced for the NICOLA cohort were funded by Economic and Social Research Council - Award Reference ES/L008459/1. CG, EB and DA are supported by SFI Award15/IA/3152, the US Ireland R&D partnership and a strategic grant from the JDRF. This study was supported, in part, by the George M. O’Brien Michigan Kidney Translational Core Center, funded by NIH/NIDDK grant 2P30-DK-081943 and by the Intramural Research Program of NIDDK. None of the funding bodies had a role in the study design or analysis.

## Declaration of interest statement/Disclosure statement

The authors declare that they have no completing interests.

## Ethics

Research ethics approval was obtained from the South and West Multicentre Research Ethics Committee (MREC/98/6/71). All participants provided written informed consent for research. No individual level data is being reported in this manuscript.

**Figures with the corresponding legend below each one**

All Supplementary Figures are provided in the separate MS Word document

## Supplementary Figures

**Supplementary Figure 1.**
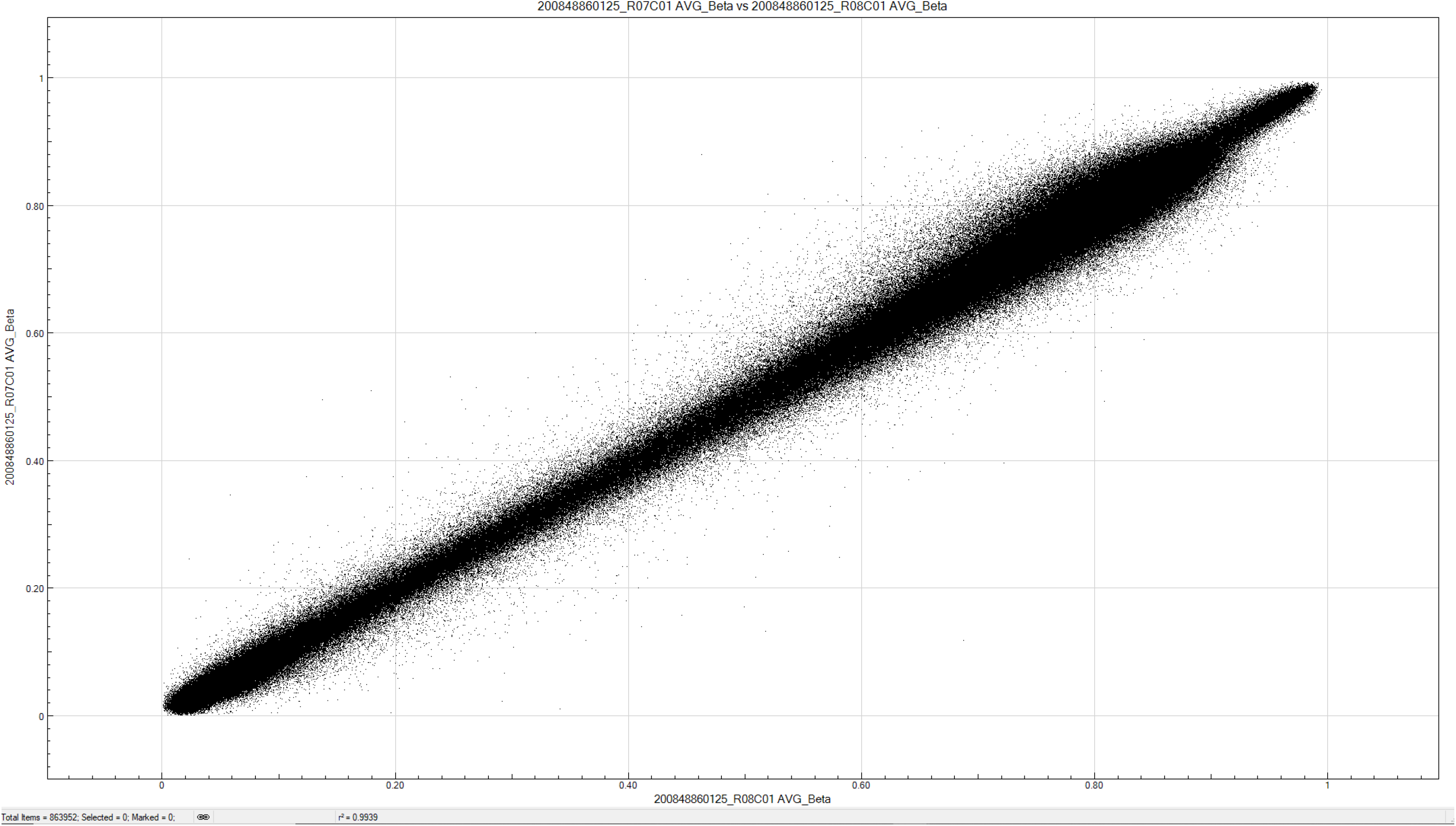
Representative concordance plots for a duplicate sample pair - average r^2^ for seven duplicates = 0.99.

**Supplementary Figure 2.**
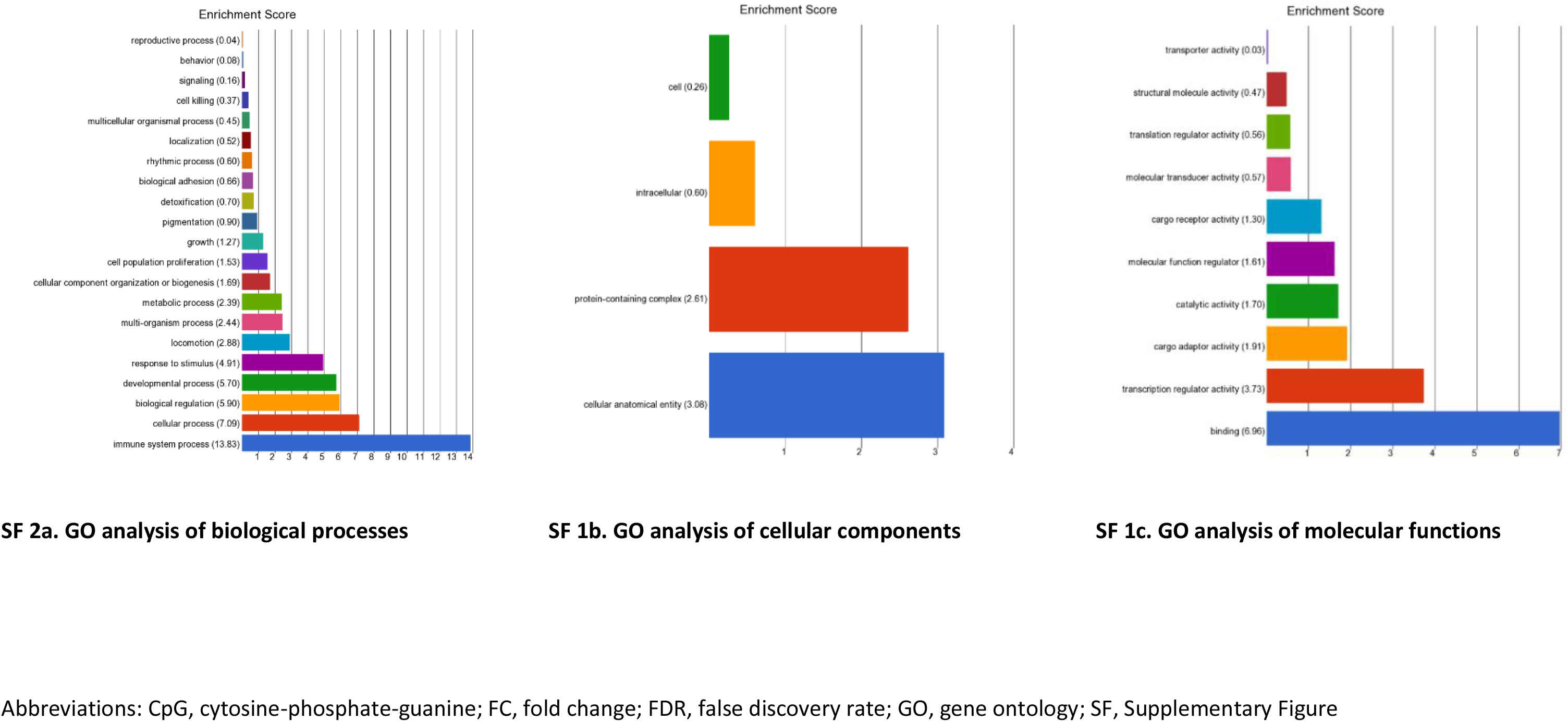
GO enrichment results for *Analysis 1*. Enrichment of top-ranked CpG sites for matched individuals with T1DM-ESKD (n=107) vs. T1DM (n=107): (FDR p-value ≤×10^−8^ and FC ≥±2). These gene classes are over-represented in the disease phenotype compared to a control gene set.

**Supplementary Figure 3.**
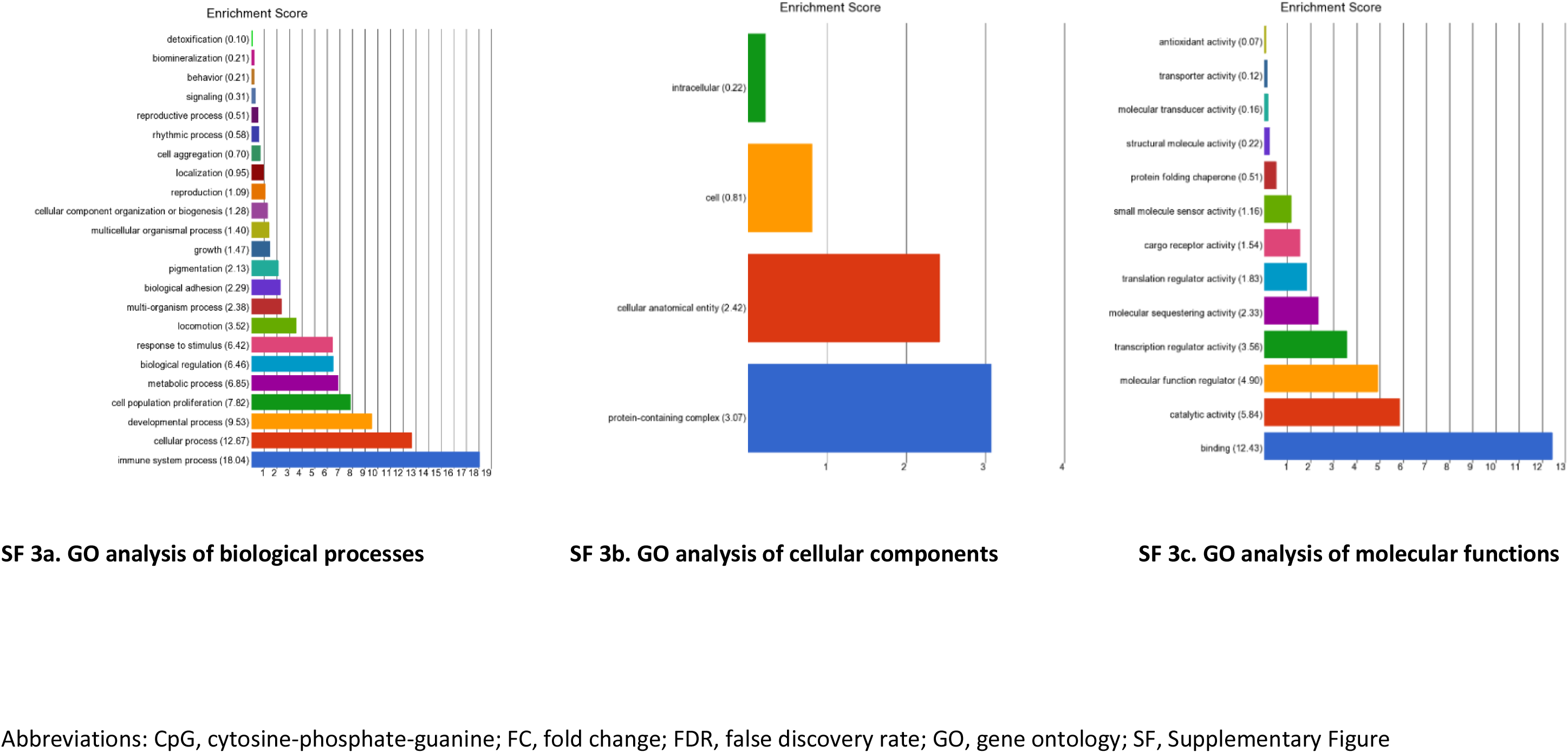
GO enrichment results for *Analysis 2*. Enrichment of top-ranked CpG sites for matched individuals with T1DM-ESKD (n=107) vs. T1DM (n=253): (FDR p-value ≤×10^−8^ and FC ≥±2). These gene classes are over-represented in the disease phenotype compared to a control gene set.

**Supplementary Figure 4.**
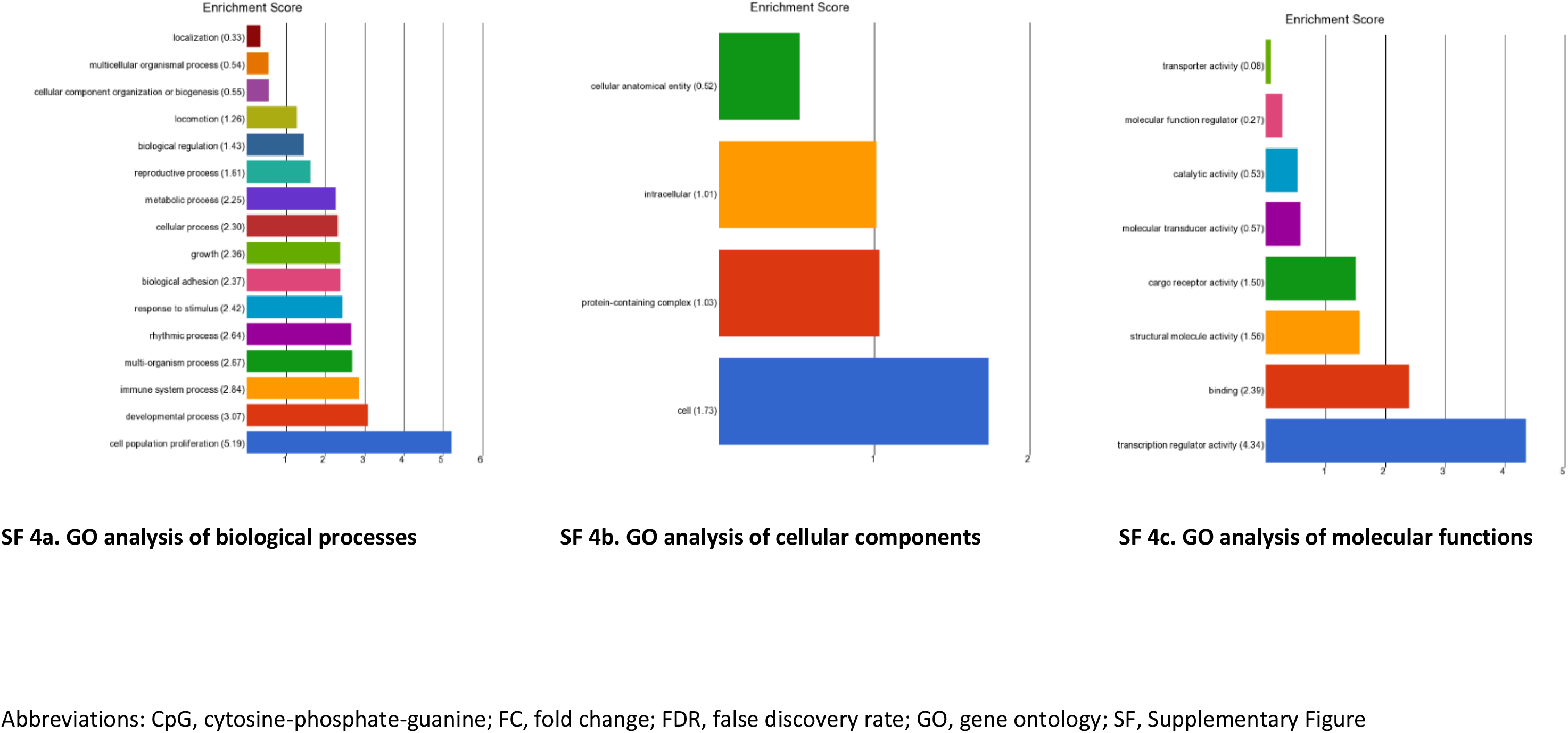
GO enrichment results for *Analysis 3*. Enrichment of top-ranked CpG sites for matched individuals with T1DM-ESKD (n=73) vs. T1DM (n=73): (FDR p-value ≤×10^−8^ and FC ≥±2). These gene classes are over-represented in the disease phenotype compared to a control gene set.

**Supplementary Figure 5.**
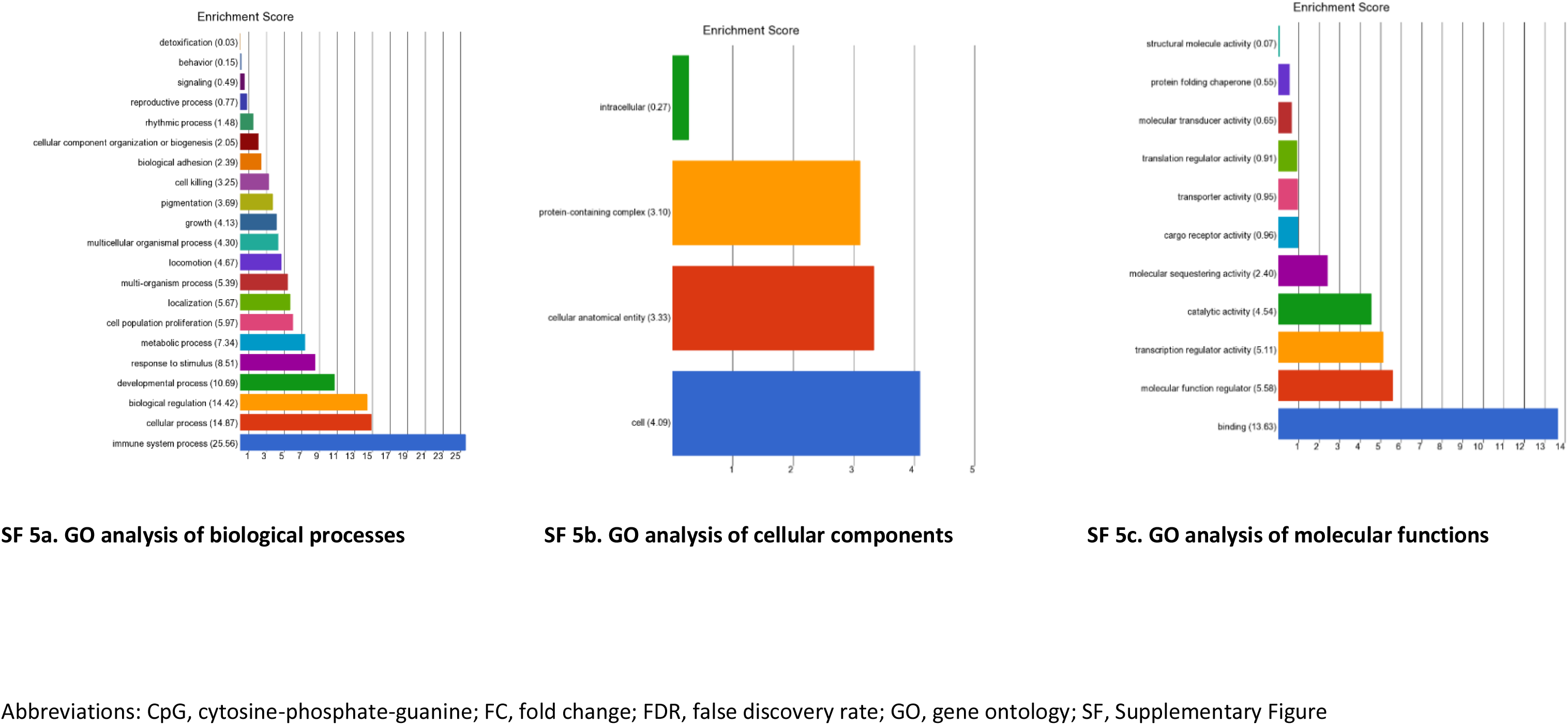
GO enrichment results for *Analysis 4*. Enrichment of top-ranked CpG sites for matched individuals with T1DM-ESKD (n=73) vs. T1DM (n=253): (FDR p-value ≤×10^−8^ and FC ≥±2). These gene classes are over-represented in the disease phenotype compared to a control gene set.

**Supplementary Figure 6.**
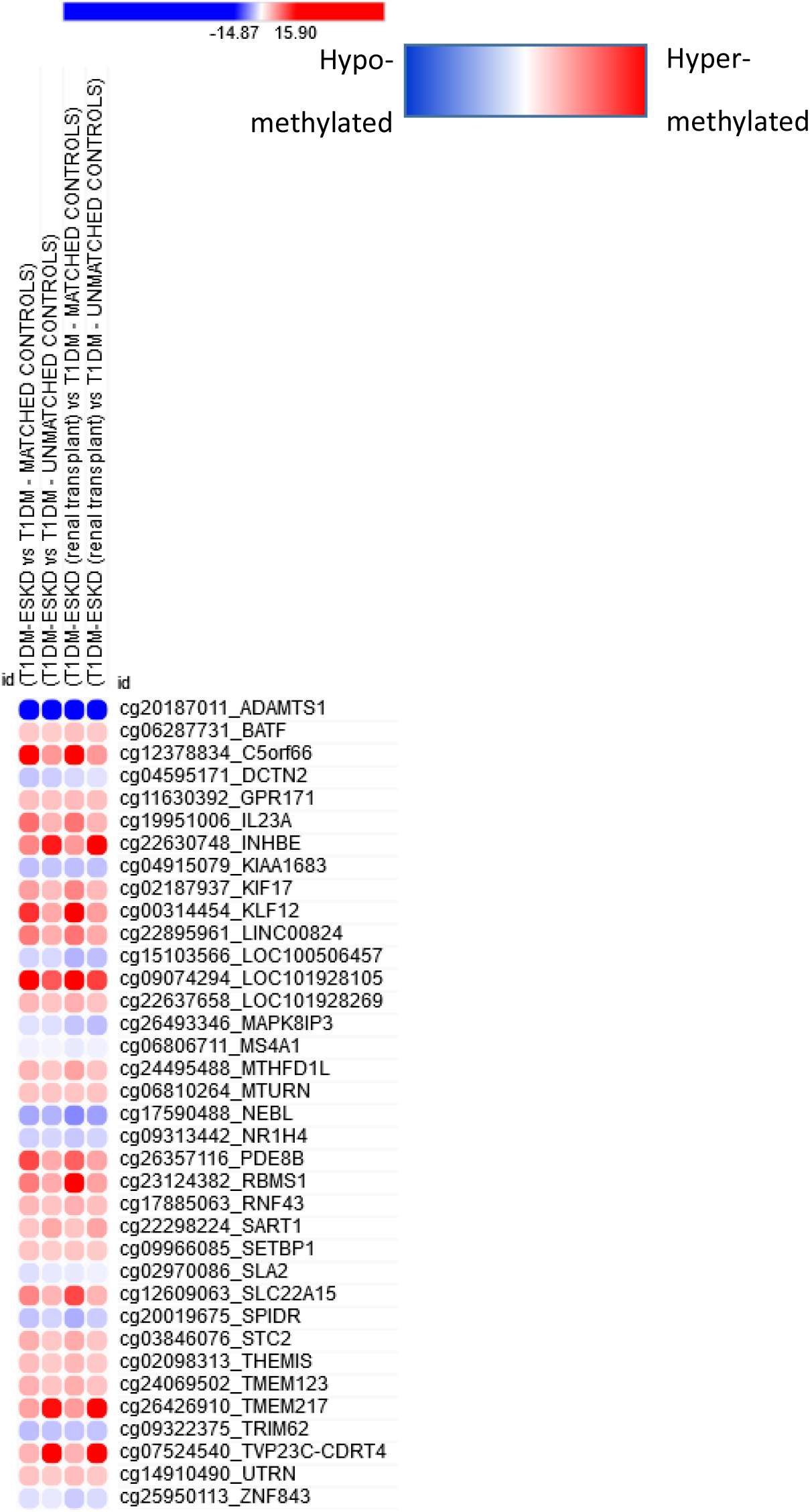
Heatmap of discovery dmCpGs in blood samples derived from T1DM ESKD versus T1DM individuals. Depicted dmCpGs are significant (p<0.05) in *Analyses 1-4.* Red = Hypermethylated; Blue = Hypomethylated.

**Supplementary Figure 7.**
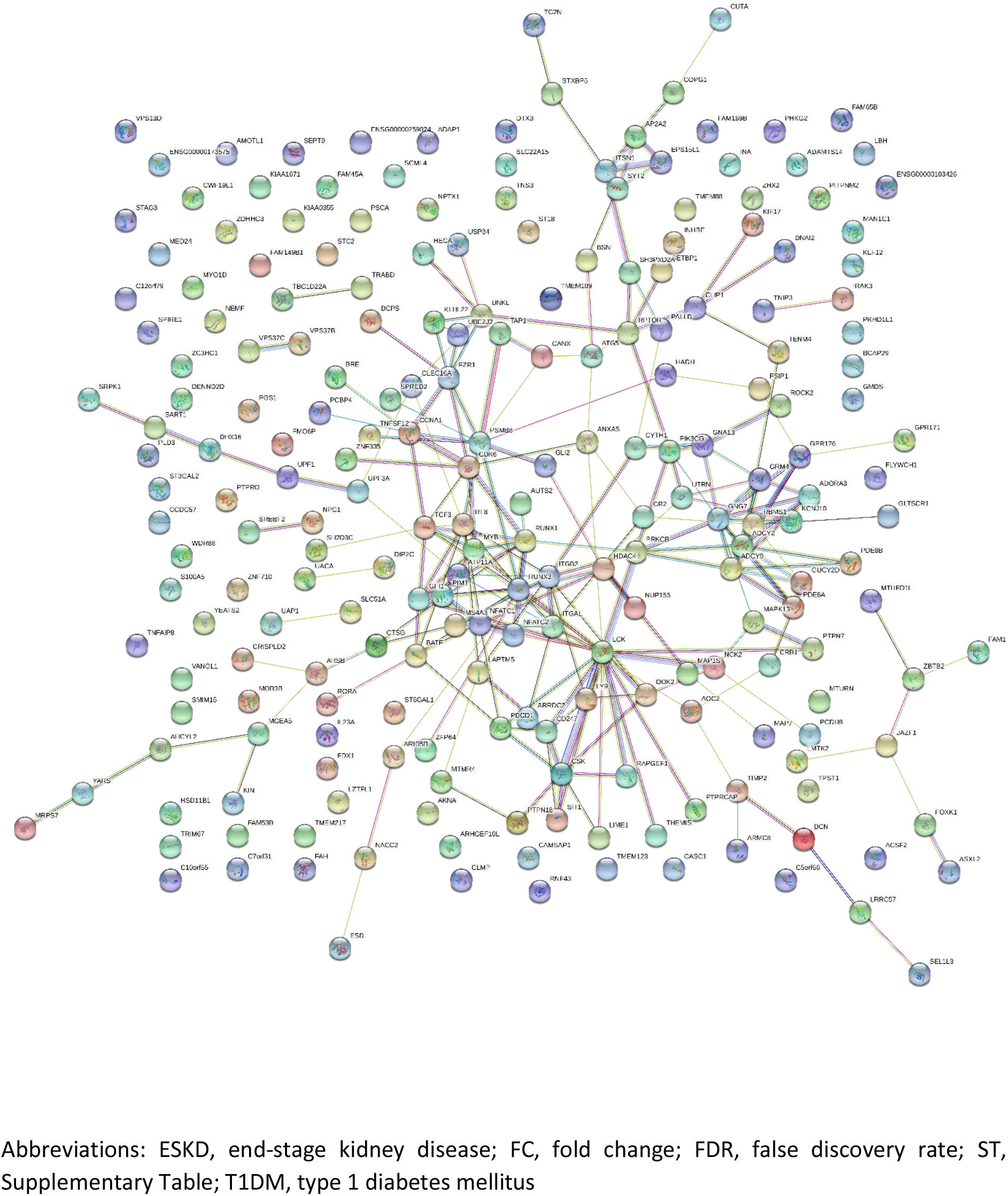
STRING analysis of genes which showed an increase in FC in individuals with T1D-ESKD from *Analysis 1*: matched individuals with T1DM-ESKD (n=107) vs. T1DM (n=107): FDR p≤×10^−8^ and FC±2 (ST4)

**Supplementary Figure 8.**
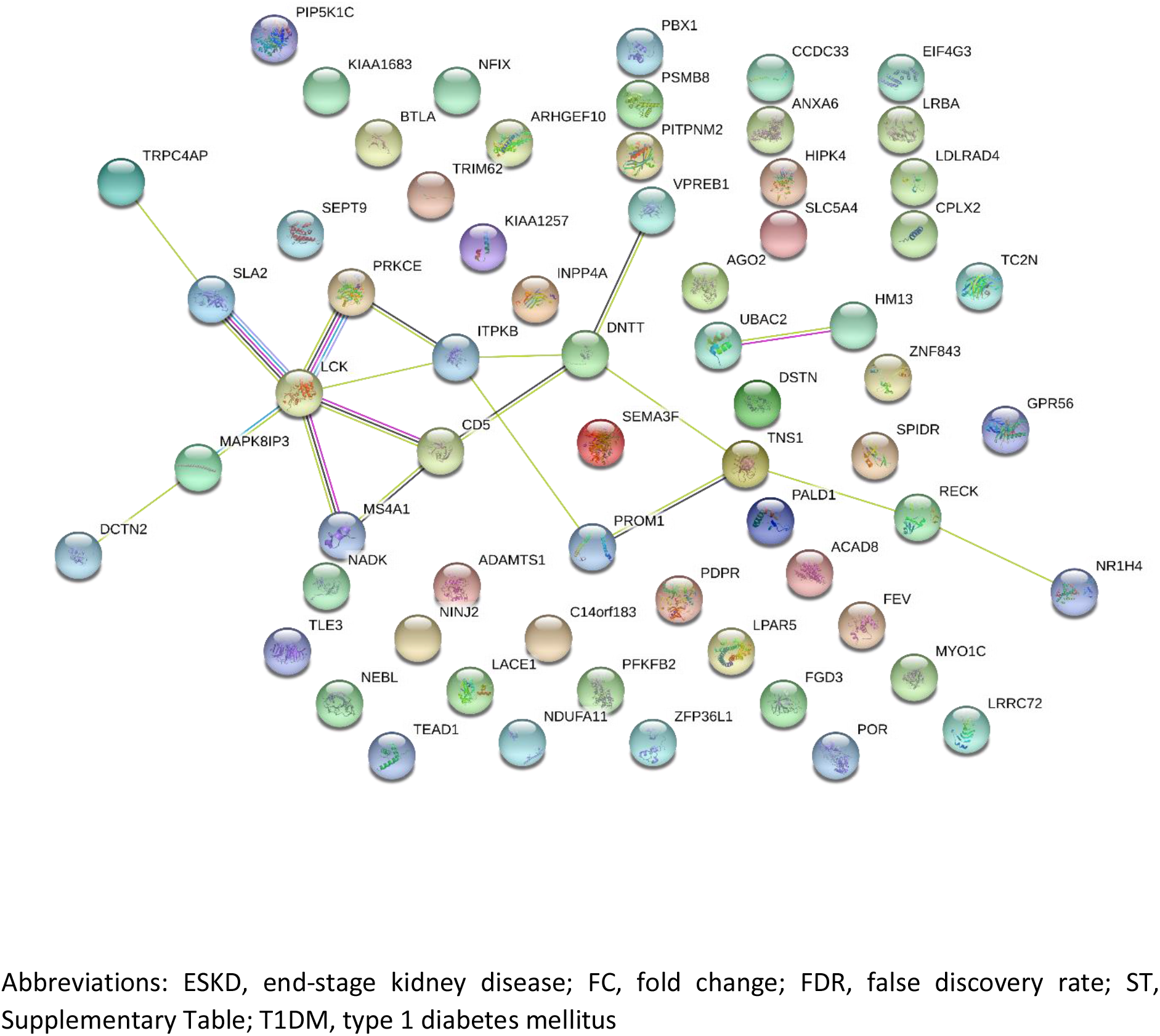
STRING analysis of genes which showed a decrease in FC in individuals with T1D-ESKD from *Analysis 1*: matched individuals with T1DM-ESKD (n=107) vs. T1DM (n=107): FDR p≤×10^−8^ and FC±2 (ST4)

**Supplementary Figure 9.**
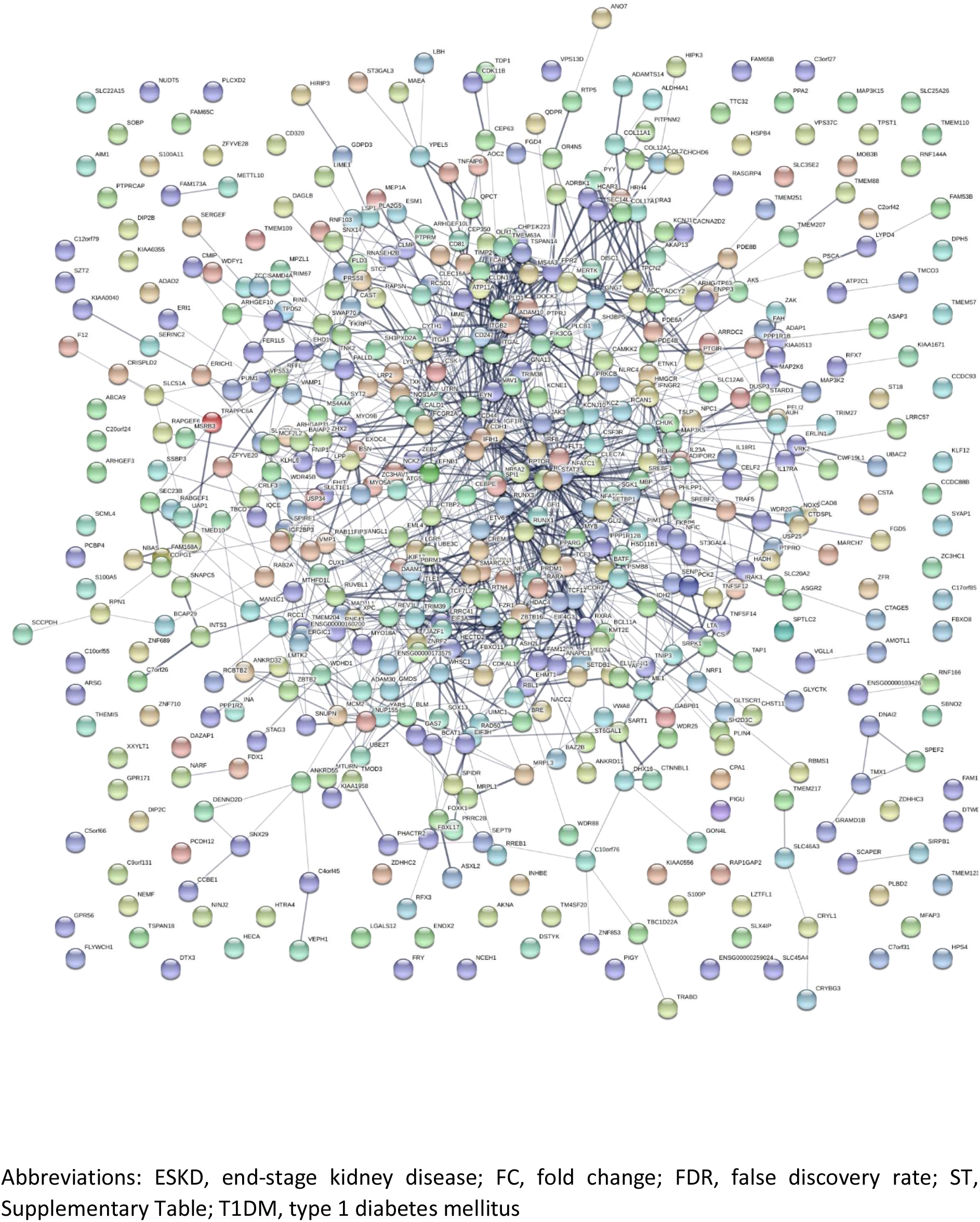
STRING analysis of genes which showed an increase in FC in individuals with T1D-ESKD from *Analysis 2*: individuals with T1DM-ESKD (n=107) vs. T1DM (n=253): FDR p≤×10^−8^ and FC±2 (ST10)

**Supplementary Figure 10.**
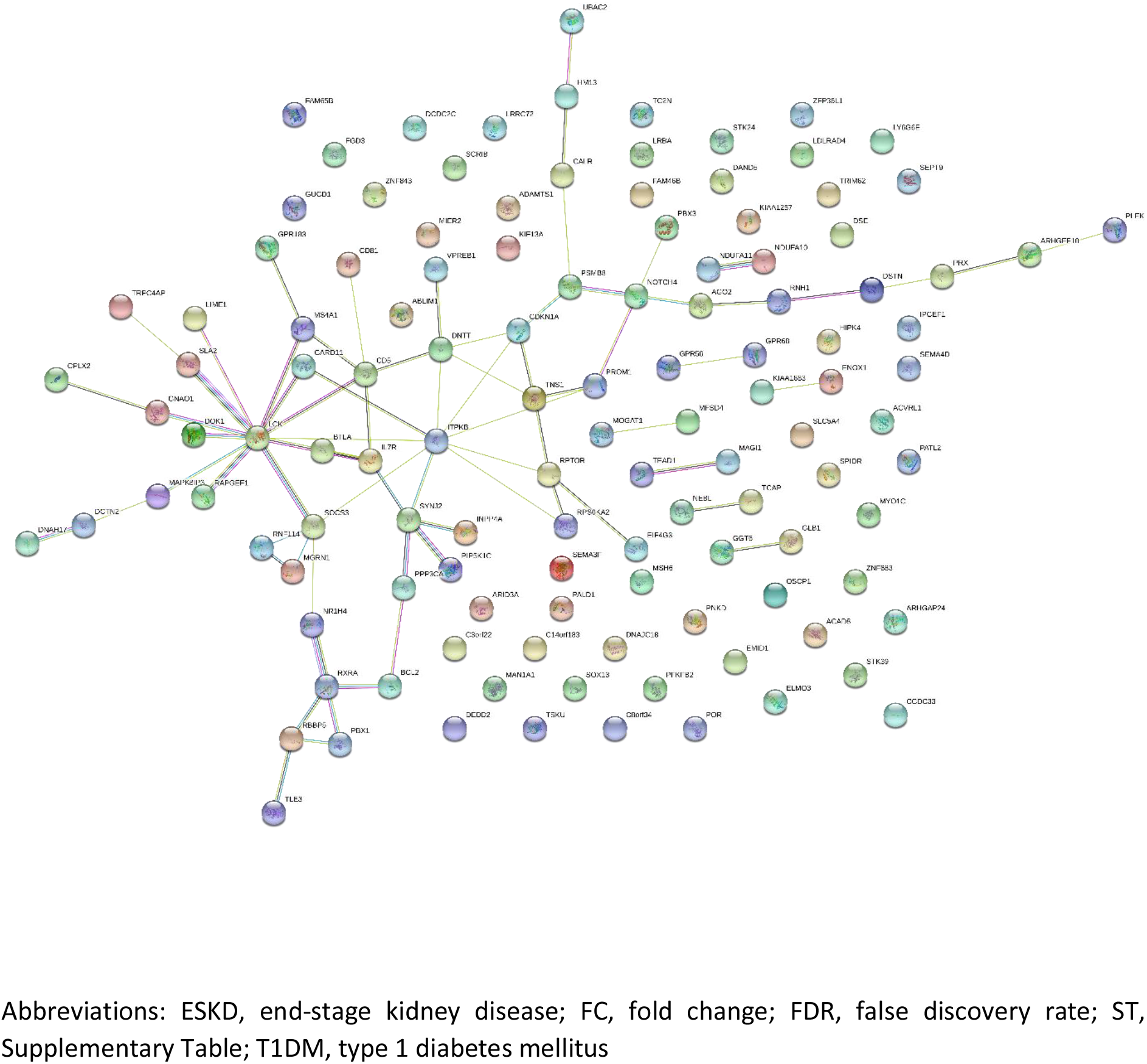
STRING analysis of genes which showed a decrease in FC in individuals with T1D-ESKD from *Analysis 2*: individuals with T1DM-ESKD (n=107) vs. T1DM (n=253): FDR p≤×10^−8^ and FC±2 (ST10)

**Supplementary Figure 11.**
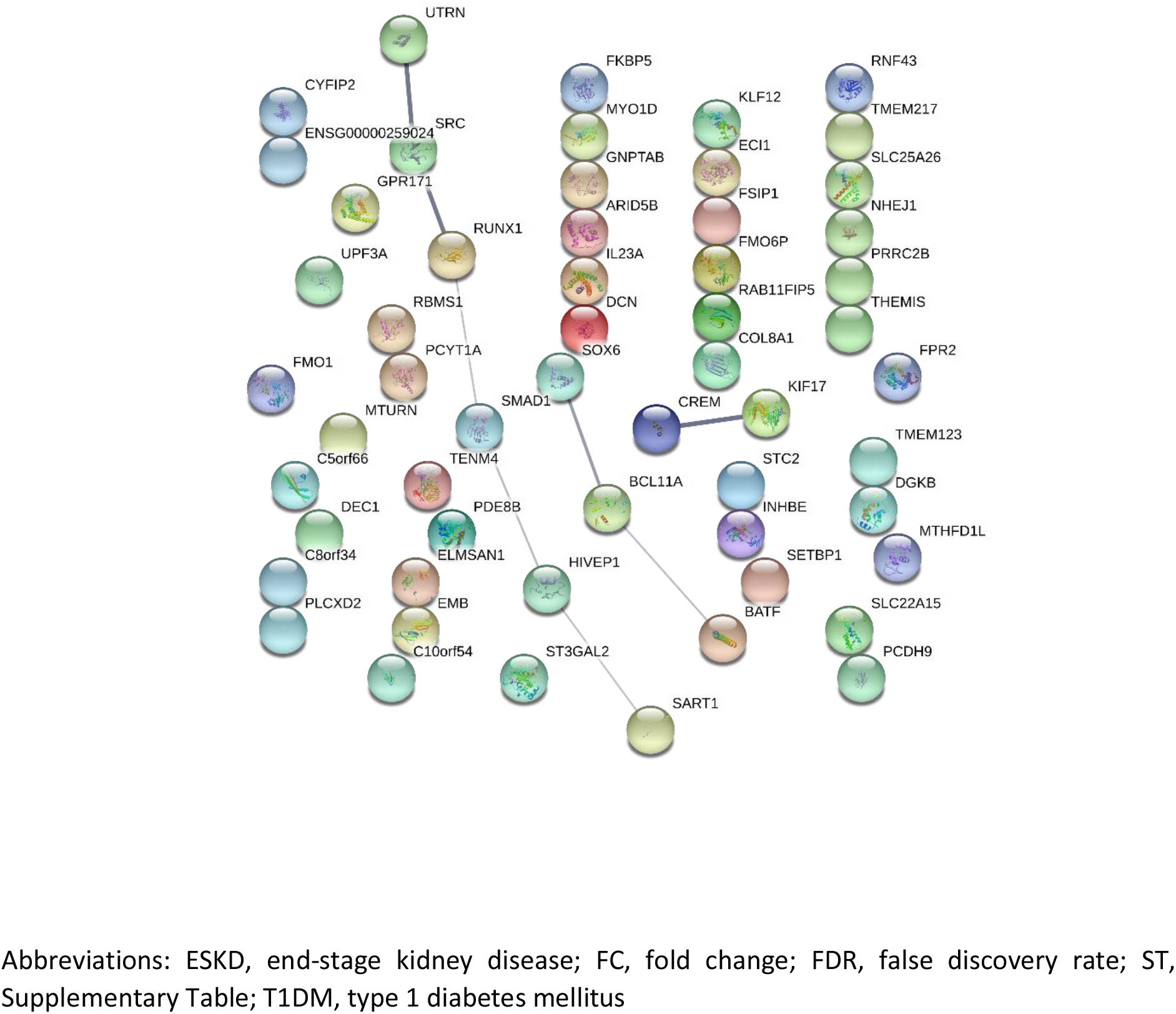
STRING analysis of genes which showed an increase in FC in individuals with T1D-ESKD from *Analysis 3*: matched individuals with T1DM-ESKD (n=73) vs. T1DM (n=73): FDR p≤×10^−^ 8 and FC±2 (ST17)

**Supplementary Figure 12.**
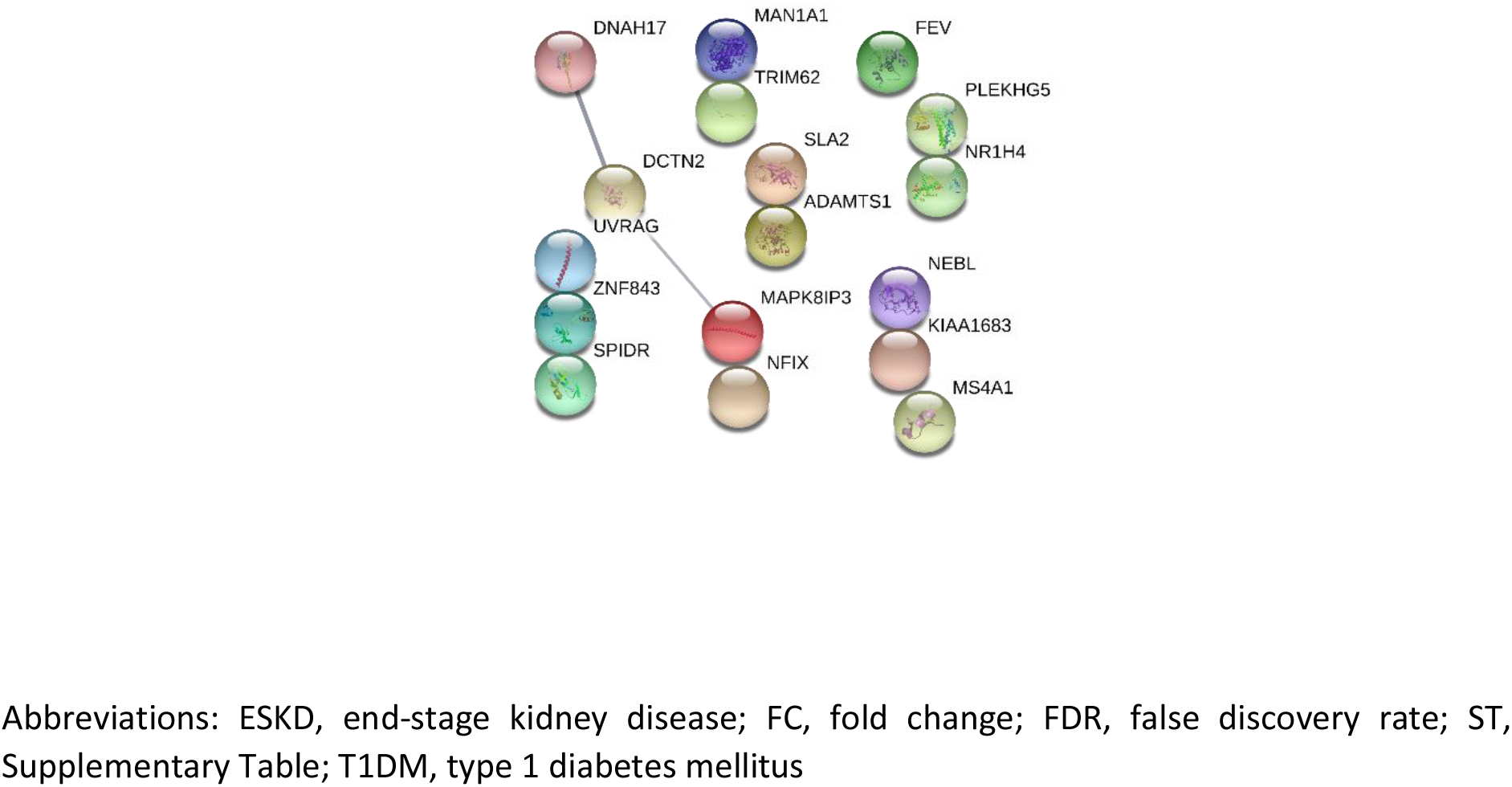
STRING analysis of genes which showed a decrease in FC in individuals with T1D-ESKD from *Analysis 3*: matched individuals with T1DM-ESKD (n=73) vs. T1DM (n=73): FDR p≤×10^−^ 8 and FC±2 (ST17)

**Supplementary Figure 13.**
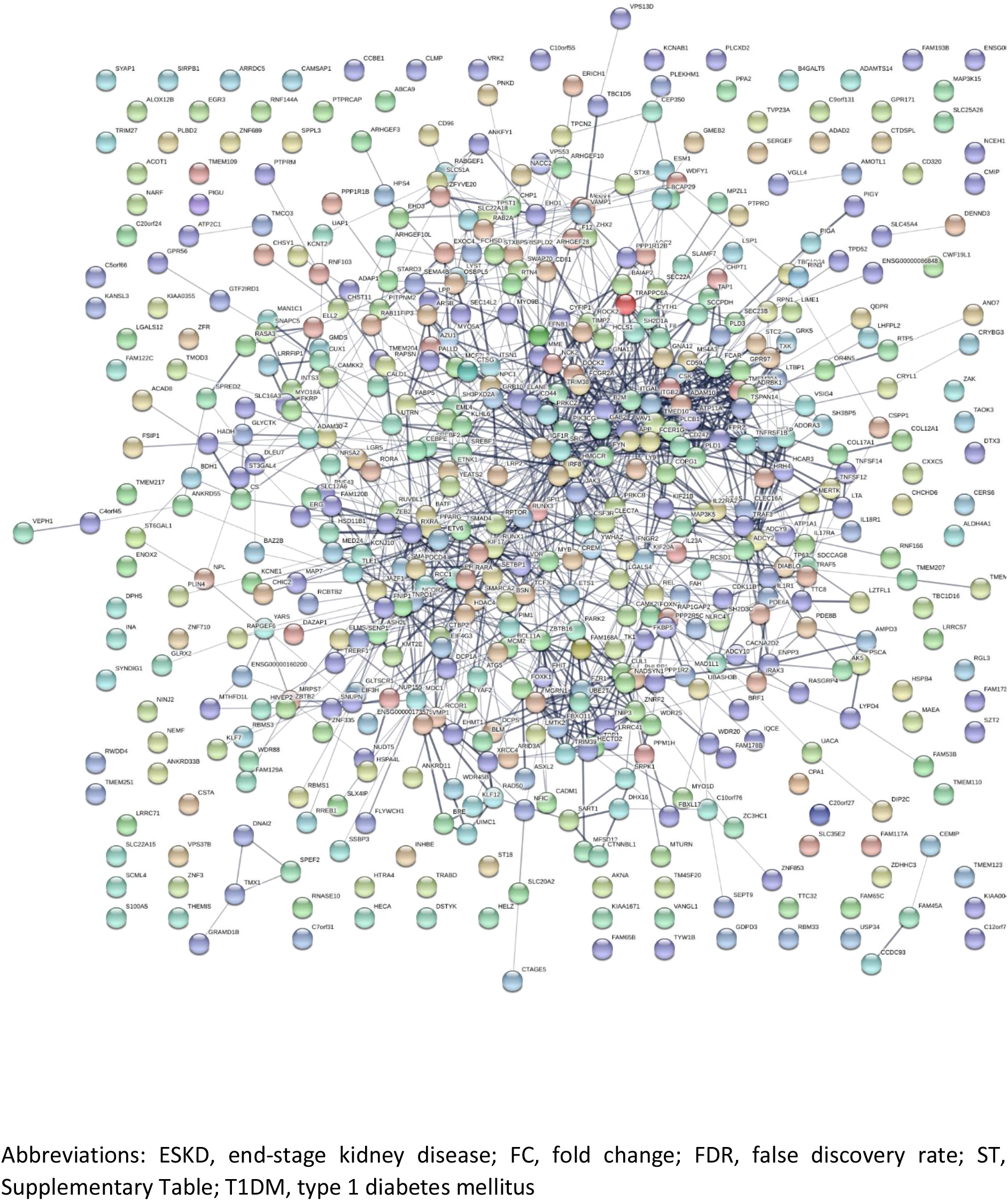
STRING analysis of genes which showed an increase in FC in individuals with T1D-ESKD from *Analysis 4*: individuals with T1DM-ESKD (n=73) vs. T1DM (n=253): FDR p≤×10^−8^ and FC±2 (ST23)

**Supplementary Figure 14.**
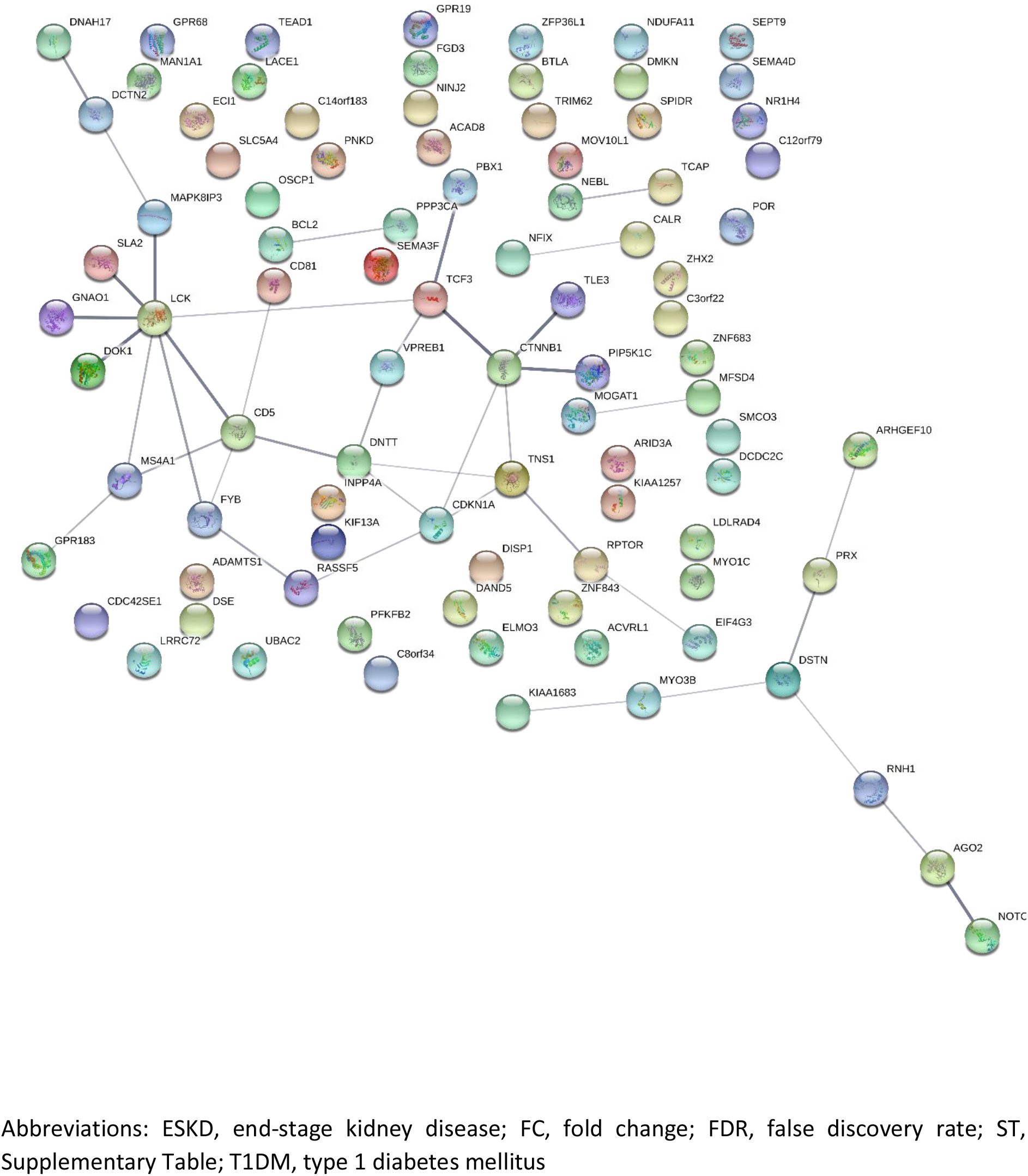
STRING analysis of genes which showed a decrease in FC in individuals with T1D-ESKD from *Analysis 4*: individuals with T1DM-ESKD (n=73) vs. T1DM (n=253): FDR p≤×10^−8^ and FC±2 (ST23)

**Supplementary Figure 15.**
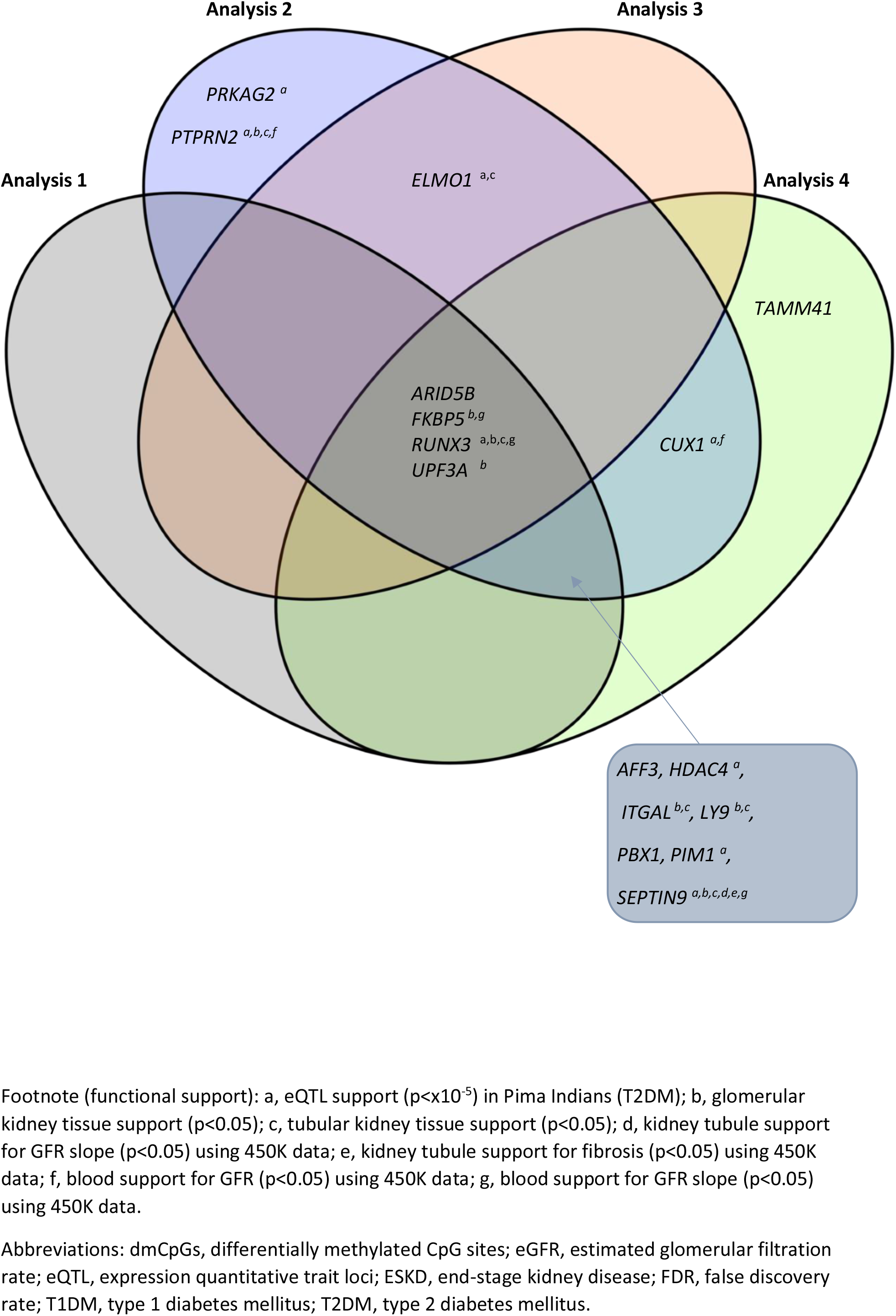
Venn diagram highlighting the genes in which top-ranked dmCpGs were located in individuals with T1DM-ESKD vs. T1DM (FDR p≤×10^−8^)

**Supplementary Figure 16.**
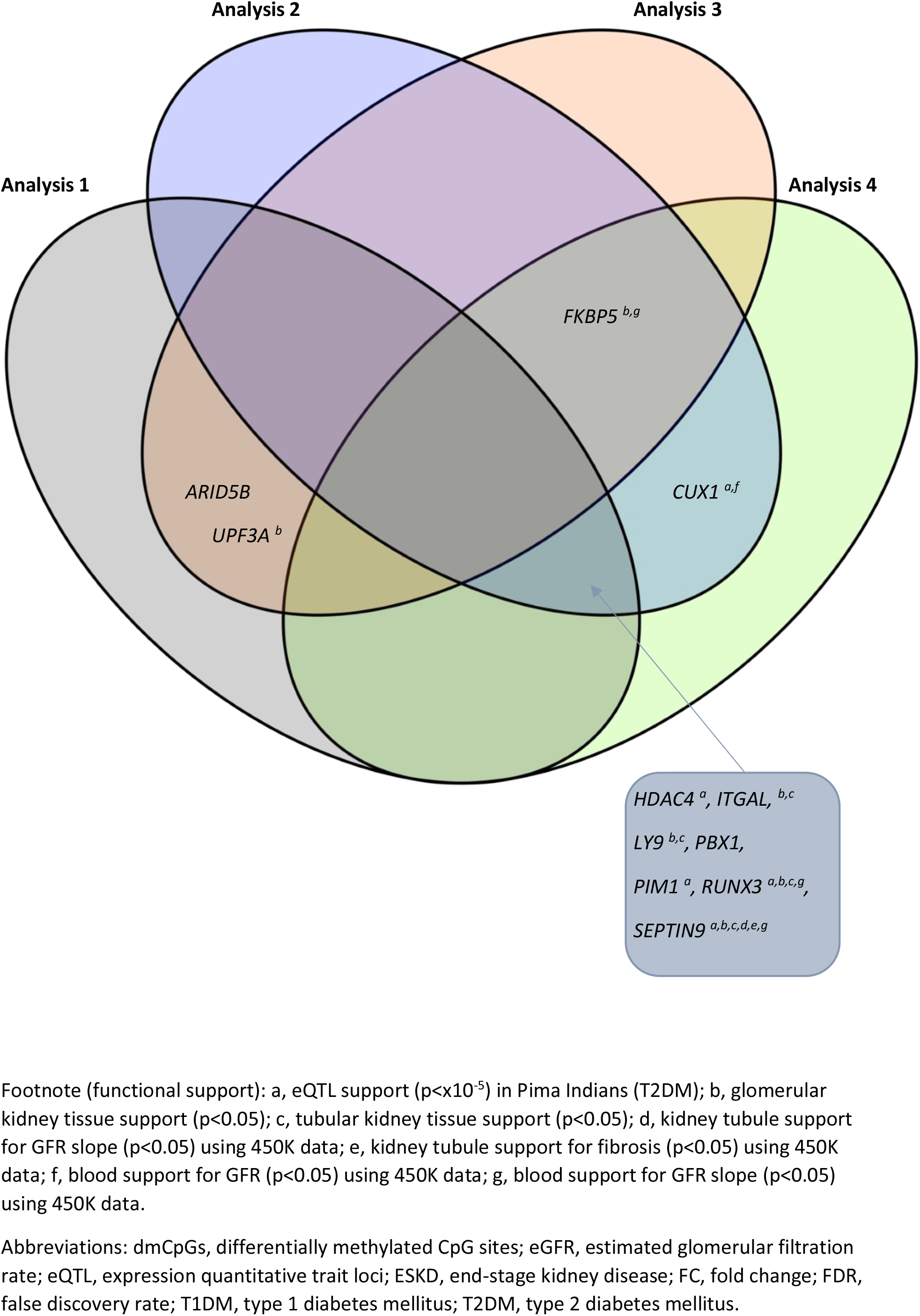
Venn diagram highlighting the genes in which top-ranked dmCpGs were located in individuals with T1DM-ESKD vs. T1DM (FDR p≤×10^−8^ and FC ±2)

## Notes

### Competing Interest Statement

The authors have declared no competing interest.

## References

1. DiMeglio LA, Evans-Molina C, Oram RA. Type 1 diabetes. Lancet. 2018;391(10138):2449–62.

2. Wang G, Ouyang J, Li S, Wang H, Lian B, Liu Z, et al. The analysis of risk factors for diabetic nephropathy progression and the construction of a prognostic database for chronic kidney diseases. J Transl Med. 2019;17(1):1–12.

3. Sandholm N, Van Zuydam N, Ahlqvist E, Juliusdottir T, Deshmukh HA, Rayner NW, et al. The genetic landscape of renal complications in type 1 diabetes. J Am Soc Nephrol. 2017;28(2):557–74.

4. UK Renal Registry (2019) UK Renal Registry 21st Annual Report. Data to 31/12/2017, Bristol, UK Available from https://www.renalreg.org/publications-reports/.

5. United States Renal Data System. 2018 USRDS annual data report: Epidemiology of kidney disease in the United States. National Institutes of Health, National Institute of Diabetes and Digestive and Kidney Diseases, Bethesda, MD, 2018.

6. Sabanayagam C, Chee ML, Banu R, Cheng CY, Lim SC, Tai ES, et al. Association of Diabetic Retinopathy and Diabetic Kidney Disease With All-Cause and Cardiovascular Mortality in a Multiethnic Asian Population. JAMA Netw open. 2019;2(3):e1915.

7. Jerram ST, Dang MN, Leslie RD. The Role of Epigenetics in Type 1 Diabetes. Curr Diab Rep. 2017;17(10):89.

8. Pociot F. Type 1 diabetes genome-wide association studies: Not to be lost in translation. Clin Transl Immunol. 2017;6(12):e162.

9. Cole JB, Florez JC. Genetics of diabetes mellitus and diabetes complications [published online ahead of print, 2020 May 12]. Nat Rev Nephrol. 2020;10.1038/s41581-020-0278-5.

10. Zhu M, Xu K, Chen Y, Gu Y, Zhang M, Luo F, et al. Identification of Novel T1D Risk Loci and Their Association With Age and Islet Function at Diagnosis in Autoantibody-Positive T1D Individuals: Based on a Two-Stage Genome-Wide Association Study. Diabetes Care. 2019;42(8):1414–21.

11. Lemos NE, Dieter C, Dorfman LE, Assmann TS, Duarte GCK, Canani LH, et al. The rs2292239 polymorphism in ERBB3 gene is associated with risk for type 1 diabetes mellitus in a Brazilian population. Gene. 2018;644:122–8.

12. Bottini N, Musumeci L, Alonso A, Rahmouni S, Nika K, Rostamkhani M, et al. A functional variant of lymphoid tyrosine phosphatase is associated with type I diabetes. Nat Genet. 2004;36(4):377–338.

13. Salem RM, Todd JN, Sandholm N, Cole JB, Chen WM, Andrews D, et al. Genome-Wide association study of diabetic kidney disease highlights biology involved in glomerular basement membrane collagen. J Am Soc Nephrol. 2019;30(10):2000–16.

14. Sandholm N, Salem RM, McKnight AJ, Brennan EP, Forsblom C, Isakova T, et al. New Susceptibility Loci Associated with Kidney Disease in Type 1 Diabetes. PLoS Genet. 2012;8(9):e1002921.

15. Pezzolesi MG, Poznik GD, Mychaleckyj JC, Paterson AD, Barati MT, Klein JB, et al. Genome-wide association scan for diabetic nephropathy susceptibility genes in type 1 diabetes. Diabetes. 2009;58(6):1403–10.

16. Sandholm N, McKnight AJ, Salem RM, Brennan EP, Forsblom C, Harjutsalo V, et al. Chromosome 2q31. 1 associates with ESRD in women with type 1 diabetes. J Am Soc Nephrol. 2013;24(10):1537–43.

17. Bansal A, Pinney SE. DNA methylation and its role in the pathogenesis of diabetes. Pediatr Diabetes. 2017;18(3):167–77.

18. Kato M, Natarajan R. Epigenetics and epigenomics in diabetic kidney disease and metabolic memory. Nat Rev Nephrol. 2019;15(6):327–45.

19. Bell CG, Teschendorff AE, Rakyan VK, Maxwell AP, Beck S, Savage DA. Genome-wide DNA methylation analysis for diabetic nephropathy in type 1 diabetes mellitus. BMC Med Genomics. 2010;3(33).

20. Smyth LJ, McKay GJ, Maxwell AP, McKnight AJ. DNA hypermethylation and DNA hypomethylation is present at different loci in chronic kidney disease. Epigenetics. 2014;9(3):366–76.

21. Gu T, Falhammar H, Gu HF, Brismar K. Epigenetic analyses of the insulin-like growth factor binding protein 1 gene in type 1 diabetes and diabetic nephropathy. Clin Epigenetics. 2014;6(1):1–6.

22. Swan EJ, Maxwell AP, Mcknight AJ. Distinct methylation patterns in genes that affect mitochondrial function are associated with kidney disease in blood-derived DNA from individuals with Type 1 diabetes. Diabet Med. 2015;32(8):1110–5.

23. Chu AY, Tin A, Schlosser P, Ko YA, Qiu C, Yao C, et al. Epigenome-wide association studies identify DNA methylation associated with kidney function. Nat Commun. 2017;8(1286).

24. Qiu C, Hanson RL, Fufaa G, Kobes S, Gluck C, Huang J, et al. Cytosine methylation predicts renal function decline in American Indians. Kidney Int. 2018;93(6):1417–31.

25. Smyth LJ, Duffy S, Maxwell AP, McKnight AJ. Genetic and epigenetic factors influencing chronic kidney disease. Am J Physiol Renal Physiol. 2014;307(7):F757–76.

26. Smyth LJ, Maxwell AP, Benson KA, Kilner J, McKay GJ, McKnight AJ. Validation of differentially methylated microRNAs identified from an epigenome-wide association study; Sanger and next generation sequencing approaches. BMC Res Notes. 2018;11:767.

27. Dirks RAM, Stunnenberg HG, Marks H. Genome-wide epigenomic profiling for biomarker discovery. Clin Epigenetics. 2016;8(122).

28. Miller SA, Dykes DD, Polesky HF. A simple salting out procedure for extracting DNA from human nucleated cells. Nucleic Acids Res. 1988;16(3):1215.

29. Ahn SJ, Costa J, Emanuel JR. PicoGreen quantitation of DNA: Effective evaluation of samples pre- or post-PCR. Nucleic Acids Res. 1996;24(13):2623–5.

30. Illumina. Infinium HD Methylation SNP List [Internet]. 2013. Available from: https://support.illumina.com/downloads/infinium_hd_methylation_snp_list.html

31. Houseman EA, Accomando WP, Koestler DC, Christensen BC, Marsit CJ, Nelson HH, et al. DNA methylation arrays as surrogate measures of cell mixture distribution. BMC Bioinformatics. 2012;8(13):86.

32. Cañadas-Garre M, Smyth LJ, Neville C, Woodside J V, Kee F, McKnight AJ. Chapter 7, Biomarkers. In: NICOLA Health Assessment Report. 2020.

33. Szklarczyk D, Gable AL, Lyon D, Junge A, Wyder S, Huerta-Cepas J, et al. STRING v11: Protein-protein association networks with increased coverage, supporting functional discovery in genome-wide experimental datasets. Nucleic Acids Res. 2019;47(D1):D607–13.

34. Weil EJ, Fufaa G, Jones LI, Lovato T, Lemley K V., Hanson RL, et al. Effect of losartan on prevention and progression of early diabetic nephropathy in American indians with type 2 diabetes. Diabetes. 2013;62(9):3224–31.

35. Berthier CC, Zhang H, Schin M, Henger A, Nelson RG, Yee B, et al. Enhanced expression of janus kinase-signal transducer and activator of transcription pathway members in human diabetic nephropathy. Diabetes. 2009;58(2):469–77.

36. Schmid H, Boucherot A, Yasuda Y, Henger A, Brunner B, Eichinger F, et al. Modular activation of nuclear factor-κB transcriptional programs in human diabetic nephropathy. Diabetes. 2006;55(11):2993–3003.

37. Fioretto P, Kim Y, Mauer M. Diabetic nephropathy as a model of reversibility of established renal lesions. Curr Opin Nephrol Hypertens. 1998;7(5):489–94.

38. Mauer M, Zinman B, Gardiner R, Suissa S, Sinaiko A, Strand T, et al. Renal and retinal effects of enalapril and losartan in type 1 diabetes. N Engl J Med. 2009;2(361):40–51.

39. Ibrahim HN, Jackson S, Connaire J, Matas A, Ney A, Najafian B, et al. Angiotensin II blockade in kidney transplant recipients. J Am Soc Nephrol. 2013;24(2):320–7.

40. Mauer M, Caramori ML, Fioretto P, Najafian B. Glomerular structural-functional relationship models of diabetic nephropathy are robust in type 1 diabetic patients. Nephrol Dial Transplant. 2015;30(6):918–23.

41. Luiza Caramori M, Kim Y, Huang C, Fish AJ, Rich SS, Miller ME, et al. Cellular basis of diabetic nephropathy: 1. Study design and renal structural-functional relationships in patients with long-standing type 1 diabetes. Diabetes. 2002;51(2):506–13.

42. Klein R, Zinman B, Gardiner R, Suissa S, Donnelly SM, Sinaiko AR, et al. The relationship of diabetic retinopathy to preclinical diabetic glomerulopathy lesions in type 1 diabetic patients: The renin-angiotensin system study. Diabetes. 2005;54(2):527–33.

43. Najafian B, Mauer M. Quantitating glomerular endothelial fenestration: An unbiased stereological approach. Am J Nephrol. 2011;33(Suppl 1):34–9.

44. Najafian B, Tøndel C, Svarstad E, Sokolovkiy A, Smith K, Mauer M. One year of enzyme replacement therapy reduces globotriaosylceramide inclusions in podocytes in Male adult patients with Fabry disease. PLoS One. 2016;11(4):e0152812.

45. Illumina. BeadArray Controls Reporter Software Guide [Internet]. 2015. Available from: https://support.illumina.com/content/dam/illumina-support/documents/documentation/chemistry_documentation/infinium_assays/infinium_hd_methylation/beadarray-controls-reporter-user-guide-1000000004009-00.pdf

46. Daca-Roszak P, Pfeifer A, Zebracka-Gala J, Rusinek D, Szybińska A, Jarzab B, et al. Impact of SNPs on methylation readouts by Illumina Infinium HumanMethylation450 BeadChip Array: Implications for comparative population studies. BMC Genomics. 2015;16(1003).

47. Nair V, Komorowsky C V., Weil EJ, Yee B, Hodgin J, Harder JL, et al. A molecular morphometric approach to diabetic kidney disease can link structure to function and outcome. Kidney Int. 2018;93(2):439–49.

48. Gluck C, Qiu C, Han SY, Palmer M, Park J, Ko YA, et al. Kidney cytosine methylation changes improve renal function decline estimation in patients with diabetic kidney disease. Nat Commun. 2019;10(2461).

49. Dang MN, Buzzetti R, Pozzilli P. Epigenetics in autoimmune diseases with focus on type 1 diabetes. Diabetes Metab Res Rev. 2013;29(1):8–18.

50. Smyth LJ, Patterson CC, Swan EJ, Maxwell AP, McKnight AJ. DNA Methylation Associated With Diabetic Kidney Disease. Frontiers (Boulder). 2020;Submitted.

51. Binder EB. The role of FKBP5, a co-chaperone of the glucocorticoid receptor in the pathogenesis and therapy of affective and anxiety disorders. Psychoneuroendocrinology. 2009;34(1):S186–95.

52. Zannas AS, Wiechmann T, Gassen NC, Binder EB. Gene-Stress-Epigenetic Regulation of FKBP5: Clinical and Translational Implications. Neuropsychopharmacology. 2016;41:261–74.

53. Zannas AS, Jia M, Hafner K, Baumert J, Wiechmann T, Pape JC, et al. Epigenetic upregulation of FKBP5 by aging and stress contributes to NF-κB-driven inflammation and cardiovascular risk. Proc Natl Acad Sci U S A. 2019;116(23):11370–9.

54. Ortiz R, Joseph JJ, Lee R, Wand GS, Golden SH. Type 2 diabetes and cardiometabolic risk may be associated with increase in DNA methylation of FKBP5. Clin Epigenetics. 2018;10(82).

55. Mihai S, Codrici E, Popescu ID, Enciu AM, Albulescu L, Necula LG, et al. Inflammation-related mechanisms in chronic kidney disease prediction, progression, and outcome. J Immunol Res. 2018;2180373.

56. Chen F, Liu X, Bai J, Pei D, Zheng J. The emerging role of RUNX3 in cancer metastasis (Review). Oncol Rep. 2016;35(3):1227–36.

57. Mei PJ, Bai J, Liu H, Li C, Wu YP, Yu ZQ, et al. RUNX3 expression is lost in glioma and its restoration causes drastic suppression of tumor invasion and migration. J Cancer Res Clin Oncol. 2011;137(12):1823–30.

58. Cen D, Xu L, Zhang S, Chen Z, Huang Y, Li Z, et al. Renal cell carcinoma: predicting RUNX3 methylation level and its consequences on survival with CT features. Eur Radiol. 2019;29(10):5415–22.

59. Wang Z, Qin G, Zhao TC. HDAC4: Mechanism of Regulations and Biological Functions. Epigenomics. 2014;6(1):139–50.

60. Hadden MJ, Advani A. Histone deacetylase inhibitors and diabetic kidney disease. Int J Mol Sci. 2018;19(9):2630.

61. Liu N, Zhuang S. Treatment of chronic kidney diseases with histone deacetylase inhibitors. Front Physiol. 2015;6(121).

62. Xiong C, Guan Y, Zhou X, Liu L, Zhuang MA, Zhang W, et al. Selective inhibition of class IIa histone deacetylases alleviates renal fibrosis. FASEB J. 2019;33(7):8249–62.

63. Boguslawska J, Kedzierska H, Poplawski P, Rybicka B, Tanski Z, Piekielko-Witkowska A. Expression of Genes Involved in Cellular Adhesion and Extracellular Matrix Remodeling Correlates with Poor Survival of Patients with Renal Cancer. J Urol. 2016;195(6):1982–902.

64. Lu Q, Ray D, Gutsch D, Richardson B. Effect of DNA methylation and chromatin structure on ITGAL expression. Blood. 2002;99(12):4503–8.

65. Parikova A, Hruba P, Krediet RT, Krejcik Z, Stranecky V, Striz I, et al. Long-term peritoneal dialysis treatment provokes activation of genes related to adaptive immunity. Physiol Res. 2019;68(5):775–83.

66. Thameem F, Wolford JK, Bogardus C, Prochazka M. Analysis of PBX1 as a candidate gene for type 2 diabetes mellitus in Pima Indians. Biochim Biophys Acta - Gene Struct Expr. 2001;1518(1–2):215–20.

67. Agha G, Hajj H, Rifas-Shiman SL, Just AC, Hivert MF, Burris HH, et al. Birth weight-for-gestational age is associated with DNA methylation at birth and in childhood. Clin Epigenetics. 2016;8(118).

68. Deucher AM, Qi Z, Yu J, George TI, Etzell JE. BCL6 expression correlates with the t(1;19) translocation in B-Lymphoblastic Leukemia. Am J Clin Pathol. 2015;143(4):547–57.

69. Tanno P Le, Breton J, Bidart M, Satre V, Harbuz R, Ray PF, et al. PBX1 haploinsufficiency leads to syndromic congenital anomalies of the kidney and urinary tract (CAKUT) in humans. J Med Genet. 2017;54(7):502–10.

70. Wei X, Yu L, Li Y. PBX1 promotes the cell proliferation via JAK2/STAT3 signaling in clear cell renal carcinoma. Biochem Biophys Res Commun. 2018;500(3):650–7.

71. Duesing K, Charpentier G, Marre M, Tichet J, Hercberg S, Balkau B, et al. Evaluating the association of common PBX1 variants with type 2 diabetes. BMC Med Genet. 2008;9(14).

72. Merkel AL, Meggers E, Ocker M. PIM1 kinase as a target for cancer therapy. Expert Opin Investig Drugs. 2012;21(4):425–36.

73. Wu Y, Deng Y, Zhu J, Duan Y, Weng WW, Wu X. Pim1 promotes cell proliferation and regulates glycolysis via interaction with MYC in ovarian cancer. Onco Targets Ther. 2018;11:6647–56.

74. Gao X, Liu X, Lu Y, Wang Y, Cao W, Liu X, et al. PIM1 is responsible for IL-6-induced breast cancer cell EMT and stemness via c-myc activation. Breast Cancer. 2019;26(5):663–71.

75. Magnuson NS, Wang Z, Ding G, Reeves R. Why target PIM1 for cancer diagnosis and treatment? Futur Oncol. 2010;9:1461–78.

76. Zhao B, Liu L, Mao J, Zhang Z, Wang Q, Li Q. PIM1 mediates epithelial-mesenchymal transition by targeting Smads and c-Myc in the nucleus and potentiates clear-cell renal-cell carcinoma oncogenesis article. Cell Death Dis. 2018;9(3):307.

77. Fu R, Xia Y, Li M, Mao R, Guo C, Zhou M, et al. Pim-1 as a Therapeutic Target in Lupus Nephritis. Arthritis Rheumatol. 2019;71(8):1308–18.

78. Neubauer K, Neubauer B, Seidl M, Zieger B. Characterization of septin expression in normal and fibrotic kidneys. Cytoskeleton. 2019;76(1):143–53.

79. Angulo JC, Andrés G, Ashour N, Sánchez-Chapado M, López JI, Ropero S. Development of Castration Resistant Prostate Cancer can be Predicted by a DNA Hypermethylation Profile. J Urol. 2016;195(3):619–26.

80. Tóth K, Galamb O, Spisák S, Wichmann B, Sipos F, Valcz G, et al. The influence of methylated septin 9 gene on RNA and protein level in colorectal cancer. Pathol Oncol Res. 2011;17(3):503–9.

81. Wu Y, Bu F, Yu H, Li W, Huang C, Meng X, et al. Methylation of Septin9 mediated by DNMT3a enhances hepatic stellate cells activation and liver fibrogenesis. Toxicol Appl Pharmacol. 2017;315:35–49.

82. Guo J, Sun C, Wang B, Ma K, Li F, Wang Y, et al. Associations between Vitamin D and β-cell function and colorectal cancer-associated tumor markers in Chinese type 2 diabetic patients with albuminuria. Clin Lab. 2019;65(4).

83. Dolat L, Hunyara JL, Bowen JR, Karasmanis EP, Elgawly M, Galkin VE, et al. Septins promote stress fiber-mediated maturation of focal adhesions and renal epithelial motility. J Cell Biol. 2014;207(2):225–35.

84. Dayeh T, Volkov P, Salö S, Hall E, Nilsson E, Olsson AH, et al. Genome-Wide DNA Methylation Analysis of Human Pancreatic Islets from Type 2 Diabetic and Non-Diabetic Donors Identifies Candidate Genes That Influence Insulin Secretion. PLoS Genet. 2014;10(3):e1004160.

85. Wang G, Watanabe M, Imai Y, Hara K, Manabe I, Maemura K, et al. Associations of variations in the MRF2/ARID5B gene with susceptibility to type 2 diabetes in the Japanese population. J Hum Genet. 2012;57:727–33.

86. Smith AR, Smith RG, Pishva E, Hannon E, Roubroeks JAY, Burrage J, et al. Parallel profiling of DNA methylation and hydroxymethylation highlights neuropathology-associated epigenetic variation in Alzheimer’s disease. Clin Epigenetics. 2019;11(52).

87. Hiwatari M, Seki M, Akahoshi S, Yoshida K, Miyano S, Shiraishi Y, et al. Molecular studies reveal MLL-MLLT10/AF10 and ARID5B-MLL gene fusions displaced in a case of infantile acute lymphoblastic leukemia with complex karyotype. Oncol Lett. 2017;14(2):2295–9.

88. Tan SH, Leong WZ, Ngoc PCT, Tan TK, Bertulfo FC, Lim MC, et al. The enhancer RNA ARIEL activates the oncogenic transcriptional program in T-cell acute lymphoblastic leukemia. Blood. 2019;134(3):239–51.

89. Xu H, Zhao X, Bhojwani D, Shuyu E, Goodings C, Zhang H, et al. ARID5B influences antimetabolite drug sensitivity and prognosis of acute lymphoblastic leukemia. Clin Cancer Res. 2020;26(1):256–64.

90. Chan WK, Bhalla AD, Le Hir H, Nguyen LS, Huang L, Gécz J, et al. A UPF3-mediated regulatory switch that maintains RNA surveillance. Nat Struct Mol Biol. 2009;16:747–53.

91. Gotoh M, Ichikawa H, Arai E, Chiku S, Sakamoto H, Fujimoto H, et al. Comprehensive exploration of novel chimeric transcripts in clear cell renal cell carcinomas using whole transcriptome analysis. Genes Chromosom Cancer. 2014;53(12):1018–32.

92. Sharma M, Brantley JG, Vassmer D, Chaturvedi G, Baas J, Vanden Heuvel GB. The homeodomain protein Cux1 interacts with Grg4 to repress p27kip1 expression during kidney development. Gene. 2009;439(1–2):87–94.

93. Livingston S, Carlton C, Sharma M, Kearns D, Baybutt R, Vanden Heuvel GB. Cux1 regulation of the cyclin kinase inhibitor p27 kip1 in polycystic kidney disease is attenuated by HDAC inhibitors. Gene X. 2019;100007.

94. Porath B, Livingston S, Andres EL, Petrie AM, Wright JC, Woo AE, et al. Cux1 promotes cell proliferation and polycystic kidney disease progression in an ADPKD mouse model. Am J Physiol - Ren Physiol. 2017;313(4):F1050–9.

95. An N, Khan S, Imgruet MK, Gurbuxani SK, Konecki SN, Burgess MR, et al. Gene dosage effect of CUX1 in amurinemodel disruptsHSC homeostasis and controls the severity and mortality of MDS. Blood. 2018;131(24):2682–97.

96. Reidy K, Kang HM, Hostetter T, Susztak K. Molecular mechanisms of Diabetic kidney disease. J Clin Invest. 2014;124(6):2333–40.

97. Ye J, Richardson TG, McArdle WL, Relton CL, Gillespie KM, Suderman M, et al. Identification of loci where DNA methylation potentially mediates genetic risk of type 1 diabetes. J Autoimmun. 2018;93(June):66–75.

98. Wu YH, Wang Y, Chen M, Zhang X, Wang D, Pan Y, et al. Association of ELMO1 gene polymorphisms with diabetic nephropathy in Chinese population. J Endocrinol Invest. 2013;36(5):298–302.

99. Turki A, Mzoughi S, Mtitaoui N, Khairallah M, Marmouch H, Hammami S, et al. Gender differences in the association of ELMO1 genetic variants with type 2 diabetes in Tunisian Arabs. J Endocrinol Invest. 2018;41(3):285–91.

100. Yahya MJ, Ismail P binti, Nordin N binti, Akim A binti M, Yusuf WS binti M, Adam NL binti, et al. Association of CCL2, CCR5, ELMO1, and IL8 Polymorphism with Diabetic Nephropathy in Malaysian Type 2 Diabetic Patients. Int J Chronic Dis. 2019;2019(2053015).

101. Peng H, Zhang Y, Zhou Z, Guo Y, Huang X, Westover KD, et al. Intergrated analysis of ELMO1, serves as a link between tumour mutation burden and epithelial-mesenchymal transition in hepatocellular carcinoma. EBioMedicine. 2019;46:105–18.

102. Arandjelovic S, Perry JSA, Lucas CD, Penberthy KK, Kim TH, Zhou M, et al. A noncanonical role for the engulfment gene ELMO1 in neutrophils that promotes inflammatory arthritis. Nat Immunol. 2019;20(2):141–51.

103. Pirini F, Noazin S, Jahuira-Arias MH, Rodriguez-Torres S, Friess L, Michailidi C, et al. Early detection of gastric cancer using global, genome-wide and IRF4, ELMO1, CLIP4 and MSC DNA methylation in endoscopic biopsies. Oncotarget. 2017;8(24):38501–16.

104. van Zuydam NR, Ahlqvist E, Sandholm N, Deshmukh H, Rayner NW, Abdalla M, et al. A Genome-Wide Association Study of Diabetic Kidney Disease in Subjects With Type 2 Diabetes. Diabetes. 2018;67(7):1414–27.

105. Köttgen A, Pattaro C, Böger CA, Fuchsberger C, Olden M, Glazer NL, et al. Multiple new loci associated with kidney function and chronic kidney disease: The CKDGen consortium. Nat Genet. 2010;42(5):376–84.

106. Yoshida T, Kato K, Yokoi K, Oguri M, Watanabe S, Metoki N, et al. Association of genetic variants with chronic kidney disease in Japanese individuals with or without hypertension or diabetes mellitus. Exp Ther Med. 2010;1(1):137–45.

107. Lee S. The association of genetically controlled CpG methylation (cg158269415) of protein tyrosine phosphatase, receptor type N2 (PTPRN2) with childhood obesity. Sci Rep. 2019;9(1):4855.

108. Abuhatzira L, Xu H, Tahhan G, Boulougoura A, Schäffer AA, Notkins AL. Multiple microRNAs within the 14q32 cluster target the mRNAs of major type 1 diabetes autoantigens IA-2, IA-2b, and GAD65. FASEB J. 2015;29(10):4374–83.

109. Yang RM, Tao J, Zhan M, Yuan H, Wang HH, Chen SJ, et al. TAMM41 is required for heart valve differentiation via regulation of PINK-PARK2 dependent mitophagy. Cell Death Differ. 2019;26:2430–46.

110. Staudt D, Stainier D. Uncovering the Molecular and Cellular Mechanisms of Heart Development Using the Zebrafish. Annu Rev Genet. 2012;46:397–418.

111. Vajdic CM, McDonald SP, McCredie MRE, Van Leeuwen MT, Stewart JH, Law M, et al. Cancer incidence before and after kidney transplantation. J Am Med Assoc. 2006;296(23):2823–31.

112. Van Leeuwen MT, Webster AC, McCredie MRE, Stewart JH, McDonald SP, Amin J, et al. Effect of reduced immunosuppression after kidney transplant failure on risk of cancer: Population based retrospective cohort study. BMJ. 2010;340(c570).

113. Alfano G, Fontana F, Colaci E, Mori G, Cerami C, Messerotti A, et al. T-cell large granular lymphocyte leukemia in solid organ transplant recipients: case series and review of the literature. Int J Hematol. 2019;110(3):313–21.

114. Woillard JB, Kamar N, Rousseau A, Rostaing L, Marquet P, Picard N. Association of sirolimus adverse effects with m-TOR, p70S6K or Raptor polymorphisms in kidney transplant recipients. Pharmacogenet Genomics. 2012;22(10):725–32.

115. Yong WS, Hsu FM, Chen PY. Profiling genome-wide DNA methylation. Epigenetics and Chromatin. 2016;9(26).

116. Mansell G, Gorrie-Stone TJ, Bao Y, Kumari M, Schalkwyk LS, Mill J, et al. Guidance for DNA methylation studies: Statistical insights from the Illumina EPIC array. BMC Genomics. 2019;20(1):366.

